# *Candida albicans* exhibits heterogeneous and adaptive cytoprotective responses to anti-fungal compounds

**DOI:** 10.1101/2022.07.20.500774

**Authors:** V Dumeaux, S Massahi, V Bettauer, S Khurdia, ACBP Costa, RP Omran, S Simpson, JL Xie, M Whiteway, J Berman, MT Hallett

## Abstract

*Candida albicans* is an opportunistic human pathogen which represents a significant threat to human health and is associated with substantial socio-economic burden. Current antifungal treatments fail at least in part because *C. albicans* can initiate a strong drug tolerance response, allowing cells to grow at concentrations above their minimal inhibitory concentration. Our goal is to better characterize this cytoprotective tolerance program at the molecular single cell level. We present here a nano-liter droplet-based fungal single cell transcriptomics platform capable of profiling thousands of individual *C. albicans* SC5314 cells in an efficient manner. Profiles of untreated cells partition into three transcriptional clusters with each highlighting a cell cycle checkpoint coupled with specific metabolic and stress responses, as perhaps expected. After just two days post-treatment with fluconazole, surviving cells bifurcate into two distinct subpopulations: the so-called α response involving upregulation of protein translation, rRNA processing and mitochondrial cellular respiration, and the β response involving processes and stress responses that assist damaged cells. By extending our time series to six days and profiling with other antifungals and bioactive compounds, we provide evidence that surviving cells transition from the α to β responses mediated by the Ribosome Assembly Stress Response (RASTR).

## Introduction

*Candida albicans* is an opportunistic human pathogen which represents a significant threat to human health (Kullberg and Arendrup, 2015; Pappas et al., 2018) and is associated with substantial socio-economic burden (Benedict et al., 2019). The fungus is the second most common cause of infectious-related deaths in extremely premature infants, and is the fourth most common cause of nosocomial bloodstream infections with a 15-20% mortality rate (Benjamin et al., 2010; Brown and Netea, 2012; Pfaller et al., 2019). Treatment with currently available antifungals can often fail due to tolerance, resistance, and clinical persistence (Delarze and Sanglard, 2015; Wuyts et al., 2018).

A population of cells is said to be *resistant* if it can grow well in the presence of an antifungal drug. Resistance is conferred by mutations that directly affect the ability of the drug to interact with its target (Berman and Krysan, 2020). A subpopulation of cells is said to be *tolerant* to a drug if it can grow at concentrations above its minimal inhibitory concentration (MIC). Drug tolerance in *C. albicans* is thought to be rooted in *phenotypic heterogeneity*, defined as the cell-to-cell variation in phenotypic response within an isogenic population. In microbes, phenotypic heterogeneity can arise through mechanisms that include stochastic or periodic expression, cell age, cell-cell interactions, chromatin modification, and genomic neoplasticity; the benefits conferred by heterogeneity in microbial populations include bet hedging, quick metabolic shifts, division of labour, and resource sharing (Ackermann, 2015). This phenotypic heterogeneity enables a subset of *C. albicans* individuals to grow slowly at otherwise inhibitory concentrations of an antifungal compound (Yu et al., 2022).

Previous work in *C. albicans* has delineated many of the core properties of tolerance including, for example, that tolerance is largely independent of drug concentration, high tolerance correlates with poor clinical outcome (Astvad et al., 2018; Levinson et al., 2021; Rosenberg et al., 2018), high tolerance increases population size (Cowen, 2005; Vincent et al., 2016), and tolerance arises too quickly to be explained by adaptive evolution (Wertheimer et al., 2016). Furthermore, adjuvants that inhibit tolerance to fluconazole (FCZ) kill *C. albicans*, thereby eliminating the evolution of resistance (Cowen, 2005; Karababa et al., 2006; Rosenberg et al., 2018; Sanglard et al., 2003b; Vincent et al., 2016).

At the molecular level, several cellular processes contribute to the complex tolerance response, including: stress responses (Bensen et al., 2004; Gong et al., 2017; Komalapriya et al., 2015; Olin-Sandoval et al., 2019; Tillmann et al., 2011; Tsai et al., 2019); cell membrane biosynthesis (Hwang et al., 2017); cell wall maintenance (Bojsen et al., 2016; Garnaud et al., 2018; Yang et al., 2017); efflux pump modulation (Coste et al., 2006); membrane properties and signaling (Llopis-Torregrosa et al., 2019; Onyewu et al., 2004; Sanglard et al., 2003a; Zarnowski et al., 2018); metabolic status (Campbell et al., 2018, 2016, 2015; Miramon and Lorenz, 2017) cellular crowding (Delarue et al., 2018); and genomic neoplasticity (Selmecki et al., 2009; Todd and Selmecki, 2020; Yang et al., 2021). The mechanisms by which so many different stress responses can regulate tolerance is not understood. For example, does it involve the *C. albican*s network of heat shock proteins and their associated signaling pathways (Gong et al., 2017), or the Environmental Stress Response (ESR) (Gasch, 2007; Gasch et al., 2000), which is prominent in *S. cerevisiae* but less general in *C. albicans* (Brown et al., 2014; Enjalbert et al., 2006) (Brown et al., 2014). Overall, we still lack a definitive list of such tolerance-related processes, and we understand very little regarding the regulatory interactions between its various components. Our primary goal here is to study the phenotypic heterogeneity of antifungal drug responses, in *C. albicans* when it grows in the presence of individual antifungal compounds. Specifically, we wanted to understand the role of phenotypic heterogeneity in drug tolerance at the transcriptional level, starting with single-cell RNAsequencing profiles.

The majority of previous studies regarding drug tolerance, which range from classic growth assays to omics profiling, have treated the surviving fungal subpopulations as a homogenous whole. This is in part because the community lacked the technology to efficiently study large numbers of individual cells. Although single cell (sc) transcriptomic assays have been applied extensively to mammalian systems, their use in fungal contexts remains limited. In the model yeast *Saccharomyces cerevisiae*, there have been limited microfluidic or barcoding sc studies examining on the order of a hundred of cells (Gasch et al., 2017; Nadal-Ribelles et al., 2019a, 2019b; Urbonaite et al., 2021). Only two high-throughout (10^3^ to 10^4^ cells) fungal transcriptome studies have been conducted in *S. cerevisiae* including Jackson and colleagues who profiled ∼40K *S. cerevisiae* cells with the commercial Chromium (10X Inc.) system (Dohn et al., 2022; Jackson et al., 2020; Jariani et al., 2020). Our previous effort was the first to profile *C. albicans* with sc-technologies (Bettauer et al., 2020); in this study we extend and refine the data and analysis that was generated using our fungal nanoliter droplet-based assay (DROP-seq), modified from the original system presented by Macosko et. al. (Macosko et al., 2015). Our system addresses technical challenges that arise in the fungal setting and provides a flexible cost-effective solution.

Here, we profiled the transcriptomes of thousands of individual cells from *C. albicans* populations that were left untreated or were challenged with antifungal compounds including fluconazole, caspofungin and rapamycin for several days. The sc-transcriptomics, in combination with “bulk” DNA-sequencing and fluorescence microscopy, allowed us to deeply examine the subpopulation composition and the phenotypic heterogeneity in isogenic *C. albicans* populations. We find that drug response is highly variable even among isogenic subpopulations and cells appear to follow distinct survival strategies involving a series of different cellular stress responses. The study highlights molecular events along this trajectory with potential for future therapeutic targeting.

## Results

### 1. An optimized single cell (sc) profiling assay to explore drug tolerance in *C. albicans*

Although sc-profiling with a commercial system is feasible in *S. cerevisiae* (Jackson et al., 2020), specific aspects of fungal biology motivated us to develop a low-cost alternative tailored for fungi. We optimized removal of the cell wall, which is 1.25 thicker in *C. albicans* than in *S. cerevisiae* (Dupres et al., 2010; Ene et al., 2012; Klis et al., 2013) to better induce stable spheroplasts. We also optimized time and concentration parameters to deliver agents that fix the transcriptome (**Methods 1**, **2**). We then constructed a nanoliter droplet-based system modified from Macosko et al. (Macosko et al., 2015) using homemade components as described by Boogeshagi et al. (Booeshaghi et al., 2019) to reduce the cost of both the device and each population assay. The profiles reported here combine our preliminary effort (Bettauer et al., 2020) with two additional batches of samples with complementary data and analyses to provide better power to examine the technical and biological efficacy of our system.

*C. albicans* populations were grown in rich media alone (untreated, UT) or with an antifungal compound: fluconazole (FCZ), caspofungin (CSP), or rapamycin (RAPA) (**Figure 1A – figure supplement 1A**). Profiling was repeated at three different timepoints. UT samples were collected at the log phase of growth while treated samples were collected at days 2 and 3 post-drug exposure. To more deeply investigate the transcriptional programs of FCZ survivors during the times when tolerance is evident, we resuspended the FCZ day-3 population in fresh YPD and samples were studied at day 6 (**Figure 1A – figure supplement 1B**). The vast majority of cells were in the yeast white morphology with less than 0.2% of cells were in a filamentous morphology (hyphae or pseudohyphae).

### 2. Single-cell profiling accurately captures transcriptional signals in *C. albicans*

After processing with the DROP-seq device, samples were sequenced following the original protocol (Macosko et al., 2015; E. Macosko et al., 2015) but with microparticle concentrations and PCR cycle numbers optimized for *C. albicans* (**Methods 3**). Gene and cell quality control are challenging exercises in all sc-profiling efforts (Svensson, 2019), but the fungal setting is especially difficult given the small amount of RNA and the stressed conditions of the fungal cells post-drug treatment (Jariani et al., 2020). In general, in single cell studies, the fundamental goal is to exclude cells where too few genes are measured, and to exclude genes that are not detected in a sufficient number of cells. Here, we sought to retain individual drug-treated cells to identify important transcripts that might reveal cellular processes and stress responses involved in tolerating the drug damage.

Sc-sequence data was processed using a reference index that covers the spliced transcriptome (He et al., 2021, p.) (**Methods 4**). Cells were considered of good quality if their profiles significantly deviated from the ambient RNA pool (FDR < 0.01) (Lun et al., 2018) (**Figure 1 – figure supplement 2A**).

The pipeline identified 18,854 high quality cells across the 6 drug/timepoint conditions with an average of 1,714 cells per sample. On average 184 transcripts were identified in each cell however there is a large dispersion in the right tail representing many cells with significantly more transcripts (max. 1,984) (**Figure 1B**). On average these transcripts arise from 94 unique genes per cell again with large right tail dispersion (max. 825) (**Figure 1 – figure supplement 2B**). Since theoretical results highlight the importance of many cells over the number of identified genes per cell (Zhang et al., 2020), we reasoned that inclusion of the sparse cells (left tail) would strengthen our analyses and help us to identify large subpopulations with strong differential transcriptional programs across the different treatments. Moreover, although the gene by cell count matrix was sparse, we observed very high concordance between FCZ pseudo-bulk profiles at day 2 and day 3 (R = 0.82; **Figure 1 – figure supplement 3A**) which indicate that the assay is robustly quantifying the expression of genes across different batches.

**Figure 1.**
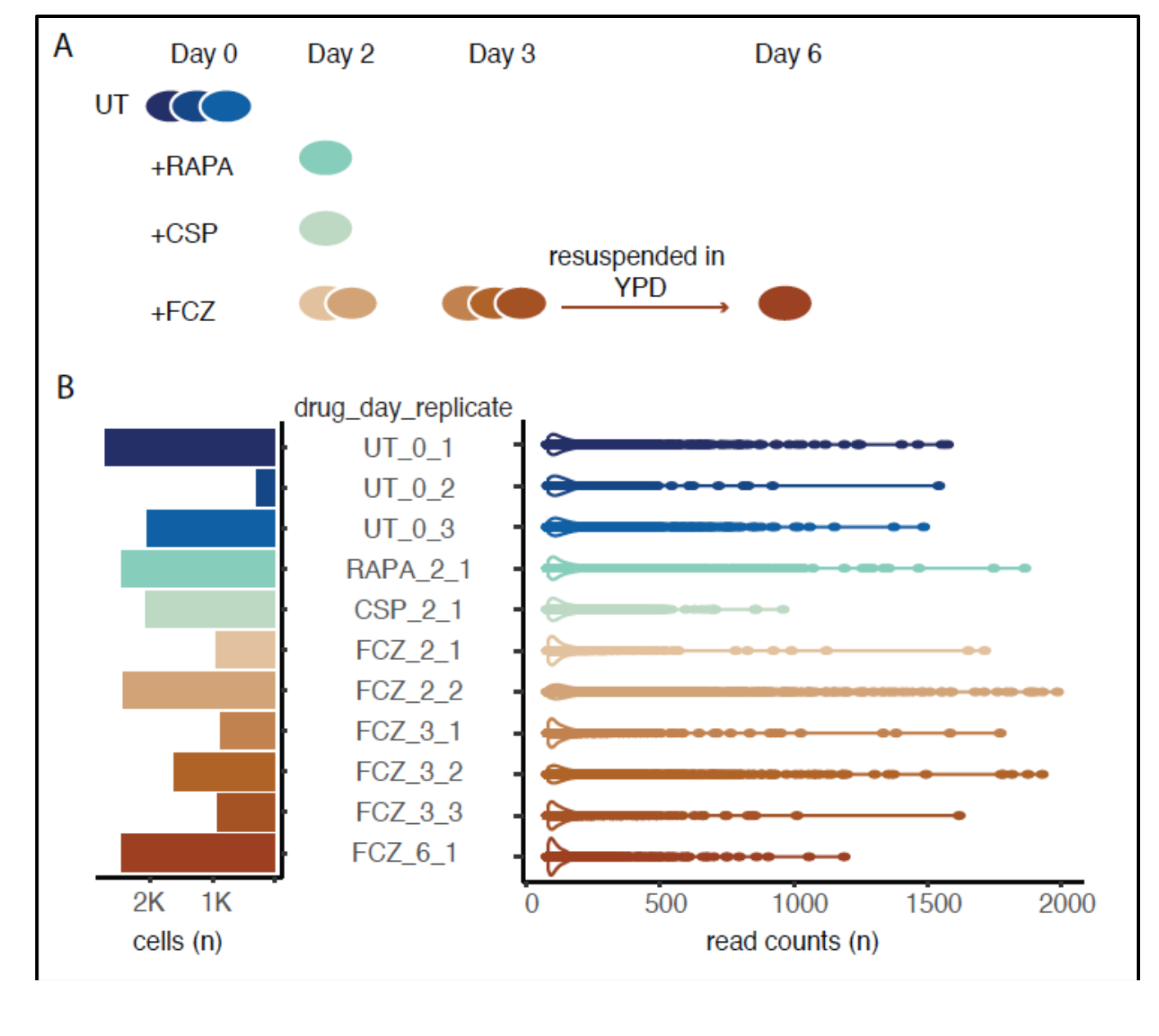
Experimental design and initial single-cell profiling. **(A)** The time series experiment begins with three replicates of untreated (UT) cells followed by profiling of rapamycin (RAPA), caspofungin (CSP) and fluconazole (FCZ, 2 replicates at day 2, and 3 replicates at day 3). After 3 days in FCZ, cells were transferred to YPD; recovered cells were profiled at day 6. **(B)** Bar plot (left) depict the number of high quality cells per sample. Violin plots (right) the distribution in the number of reads assigned to each cell.

To further investigate the robustness of the assay, we performed bulk RNA-sequencing of FCZ-treated cells at day 2 post-exposure (**Methods 5**), and compared this bulk profile with the pseudo-bulk derived from mapping single-cell reads to the reference genome but ignoring barcodes (unfiltered “pseudo-bulk” profiles; **Methods 6**). Both methods identified 6071 genes with only 172 genes not detected in one or more of the pseudo-bulk profiles. We note that missing genes were mostly expressed at low levels in the bulk profile (**Figure 1 – figure supplement 3B**). Moreover, day 2 and day 3 pseudo-bulk replicates were significantly correlated with the bulk RNA-sequencing (**Figure 1 – figure supplement 3C**). This strongly suggests that the DROP-seq-derived profiles sample the *C. albicans* transcriptome, capturing true biological signals, primarily missing transcripts expressed at lower levels.

### 3. Heterogeneous transcriptional profiles highlight metabolic and stress responses coupled with cell cycle checkpoints in isogenic untreated cells

To identify major sources of cell-to-cell variability in isogenic untreated *C. albicans* populations, mid-log phase cells grown under standard conditions were collected for sc-profiling. In addition to sc-transcriptomes, “bulk” DNA-sequencing profiles were generated to verify strain isogenicity (**Methods 7**).

To identify the main sources of cell-to-cell variability in UT cells, we applied an unsupervised dimension reduction approach, clustered the sc-transcriptional data and then visualized the data consisting of N=4,877 UT cells with a two dimensional UMAP (**Methods 8**). This analysis provides preliminary evidence for the existence of three clusters of cells with different gene expression patterns (**Figure 2A**). To understand possible biological features that distinguish these three clusters, we used prior transcriptional profiling studies to curate 43 gene signatures of processes likely to be involved in microbial phenotypic heterogeneity and drug tolerance. These included: the cell cycle, specific and general stress responses, the TCA cycle, glycolysis and other metabolic pathways, as wells as efflux pumps/transporters, ergosterol biosynthesis and other genes coding for essential components of the fungal cell membrane (**Figure 2 – table supplement 1, Methods 8**). In most cases, these expression signatures used bulk transcriptional studies, either in *C. albicans* directly or in other fungi; where necessary, we identified orthologues of relevant genes in *C. albicans* (Balakrishnan et al., 2012), while taking into account regulatory re-wiring between these species (Johnson, 2017; Lavoie et al., 2010).

**Figure 2.**
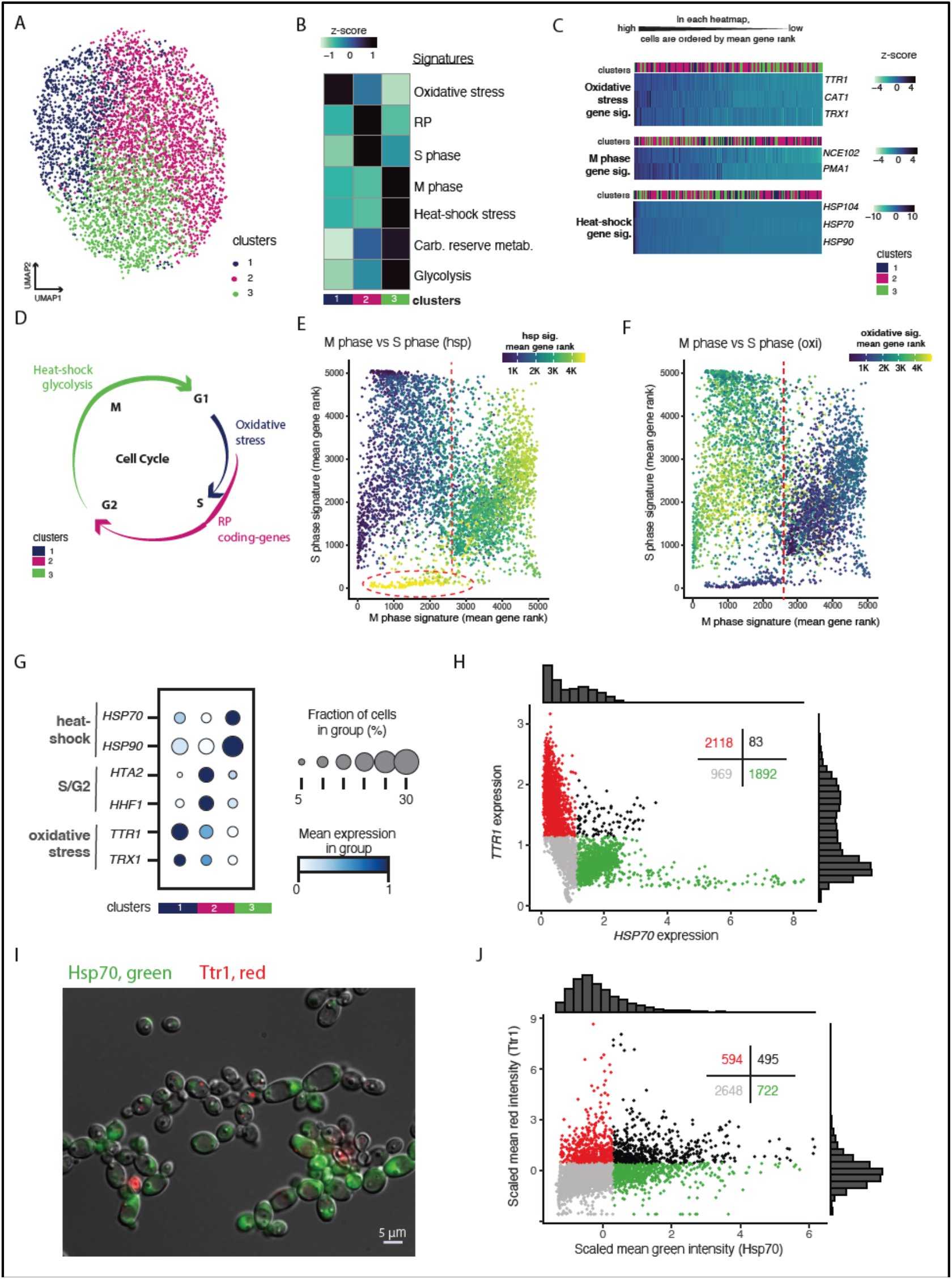
Cell to cell heterogeneity in untreated (UT) cell populations. **(A)** UMAP embedding of UT cells. Leiden clustering identified 3 cell clusters. **(B)** Average expression levels (VISION z-score, color bar) of curated signatures across the three clusters. **(C)** For each signature, the heatmap depicts the expression of selected genes (rows) across cells (columns). We used MAGIC to impute expression levels (z-score, color bar). Cells are linearly ordered by the overall magnitude of gene expression (mean gene rank). “Clusters” (top) indicate in which cluster the cell is assigned. **(D)** Simplified schematic of the cell cycle; colors correspond to the three clusters in which heat-shock and glycolysis, oxidative stress, and ribosomal protein-coding (RP) signature were most active. **(E, F)** For each cell (point) in the scatterplots, we are comparing the cells’ overall magnitude of gene expression (mean gene rank) of the M and S phase signatures. Color bars indicate the cells’ overall magnitude of gene expression of the **(E)** heat-shock or **(F)** oxidative stress signature. **(G)** Expression of selected genes that are markers for distinct clusters and which belong to signatures relevant to our analysis. Color-bar indicates the mean expression across cells in each cluster. The size of the dot indicates the proportion of cells that express the gene within each cluster. **(H)** Scatterplot of cells (dots) based on expression level imputed using MAGIC. Colors indicate: red, *TTR* expression >1.2, *HSP70* expression <1.2 ; green, *TTR* expression <1.2, *HSP70* expression >1.2; black, both *TTR1* and *HSP70* expression >1.2; and grey, expression of both genes was <1.2. Distributions of expression are illustrated in histograms above and to the right and number of cells in each group is provided in the top right of the figure. **(I)** Representative microscopy image of RFP-tagged TTR1 and GFP-tagged HSP70 cells **(J)** Plot of the mean intensities captured in microscopy images of RFP-tagged TTR1 and GFP-tagged HSP70.

The strength of each signature was measured in each cell using the sc-expression profiles (**Methods 8**). The most variable signatures across the three clusters of UT cells are displayed in **Figure 2B**, where color corresponds to the average score (z-score, colorbar) across all cells within each cluster. A complementary, more detailed perspective of the most variable genes in these signatures is found in **Figure 2C**, where cells are ordered by the overall magnitude of expression of the selected genes as captured by the mean gene rank (**Methods 8)**. Notably, cells in cluster 1 (blue) are associated with oxidative stress; cluster 2 (darkpink) is associated with the S phase and ribosomal proteins (RP) (see also **Figure 2 - figure supplement 1A-C**), and cluster 3 (green) is associated with the M phase, glycolysis, carbohydrate reserve metabolism, and heat-shock stress genes. The relationships between stress responses, metabolic processes and cell cycle are summarized in **Figure 2D**.

Since several previous reports associated cell cycle phase with the expression of genes involved in stress responses and metabolism (Brauer et al., 2008; Chiu et al., 2011; Hossain et al., 2021; Senn et al., 2012), we further investigate the patterns of co-expression of these processes in individual cells. First, we re-affirmed that cells with higher expression of genes involved in M phase also have high expression of the heat-shock response (**Figure 2E**). This is consistent with a previous report demonstrating the role of heat-shock proteins in the regulation of mitotic exit (Senn et al., 2012). We also observe that the heat-shock response reaches its relative highest expression in a small group of cells with concomitant low expression of the M and S phase, glycolysis and RP-coding genes (**Figure 2E – figure supplement 2A,B**). These few cells are likely in cell cycle arrest due to high stress levels.

Finally, **Figure 2F** shows that cells that are not in M phase have low expression of the heat-shock signature and high relative expression of the oxidative stress signature. **Figure 2F – figure supplement 2A** also establishes that non-M phase cells have low expression of genes involved in glycolysis. From this, we infer that these cells are likely early in the cell cycle and may perhaps enter a diauxic shift in response to nutrient limitations as glucose is rapidly exhausted from the growth medium (Maris et al., 2001; Uppuluri and Chaffin, 2007).

To further test the assumption that genes are expressed differently in these three cell clusters, we selected pairs of genes that the sc-transcriptomics profiles predicted would have mutually exclusive expression in any given cell (**Figure 2G**). A strong example of this involves the Heat-Shock Protein 70 (*HSP70)*, which is predominantly expressed in cluster 3 (green), and dithiol glutaredoxin (*TTR1),* which is predominantly expressed in cluster 1 (blue). These two genes had significant mutually exclusive expression patterns (**Figure 2H**; McNemar test, p-value < 0.001). Importantly, we built a dual fluorescent reporter strain with GFP-tagged HSP70 and RFP-tagged TTR1; fluorescence microscopy analysis also exhibited a highly significant degree of mutually exclusivity in expression (**Figure 2IJ – figure supplement 3,** McNemar test, p-value < 0.001).

These results support the idea that the distinct stress responses in UT population data are not due to stress responses during single-cell profiling. Rather, they likely reflect the cell cycle phase and/or the metabolic state of that cell. The microscopy results also support the existence of the clusters identified through sc-transcriptomics and highlight the importance of metabolic- and stress-sensitive steps in cell cycle progression.

In studies of bulk populations of *S. cerevisiae*, multiple specific-stress responses have been previously detected with simultaneously expression, however the existence of an analogous general stress response in *C. albicans* has been debated (Brown et al., 2020; Enjalbert et al., 2003; Gasch, 2007; Gasch et al., 2000). Single-cell profiling has the potential to determine if distinct subpopulations of cells express different stress responses as presented above. Our data suggests that cells do not express genes from the entire general induced Environmental Stress Response (iESR) previously identified in *S. cerevisiae* (**Figure 2 – figure supplement 4A**). Similarly, we did not observe concordant expression of genes of the general Hog1-driven response identified in highly stressed *C. albicans* cells (Brown et al., 2020) (**Figure 2 – figure supplement 4B**). We note that UT cells here were grown *in vitro* in the absence of any known stresses. Individual UT cells expressed only one, rather than multiple, stress response pathways and the expression of these pathways was associated with a cell cycle phase and metabolic state.

### 4. Exposure to antifungal compounds: emergence of “comets”

We next investigated the *C. albicans* response to anti-fungal compounds using fungistatic fluconazole (1 µg/mL, 1× MIC_50_), and fungicidal caspofungin (1 ng/mL, <.03x MIC). These concentrations were chosen to ensure a sufficient number of survivors (Cappelletty and Eiselstein-McKitrick, 2007; Diaz et al., 2016; Girmenia et al., 2000; Sanglard et al., 1998; Shokohi et al., 2016; Stevens et al., 2004; Yang et al., 2017; Yuksekkaya et al., 2011). Since low doses of rapamycin inhibit tolerance to FCZ (Rosenberg et al., 2018), we also profiled populations with a subinhibitory concentration of RAPA (MIC_80_ < 1 µg/mL, 0.5 ng/ml (.0005x MIC)) (Cruz et al., 2001). At these concentrations, FCZ treatment slows growth relative to UT controls in the first days after exposure, while subinhibitory doses of CSP and RAPA did not significantly affect growth as measured by OD_600_ **(Figure 3 – figure supplement 1A)**.

We focused on 11,309 good quality cells captured at 2 or 3 days post-exposure to FCZ, CSP and RAPA, in addition to the 5,062 UT cells. Unsupervised clustering of the sc-expression profiles revealed a total of 24 clusters, with the four main clusters containing 92% of all N=16,371 cells (**Figure 3A,B**). The fifth largest cluster (5-purple, N = 501 cells, 3%) lies closely adjacent to the largest cluster (1-darkpink) in the two dimensional UMAP visualization. UT cells are found primarily in cluster 3-green and to a lesser extent in cluster 2-pink (**Figure 3B**). UT-enriched regions have the highest expression of the glycolysis and carbohydrate reserve metabolism pathways (**Figure 3C – figure supplement 1B-C**). This is consistent with the fact that UT cells were collected during log-phase before glucose became a limiting factor, while all non-UT cells were harvested at least two days post-media renewal.The remaining 3% of cells (N = 573) were outliers scattered across 19 distinct clusters (others cluster-darkblue; **Figure 3A**). The clusters appear to radiate like “comets” from the center of the UMAP. This pattern suggests that the small set of cells from each comet have strong transcriptional similarity but each such comet is transcriptionally distinct from the other comets. The comets were primarily observed in FCZ survivors at 2 days (462 of 573 cells in the 19 comet clusters; **Figure 3B**).

**Figure 3.**
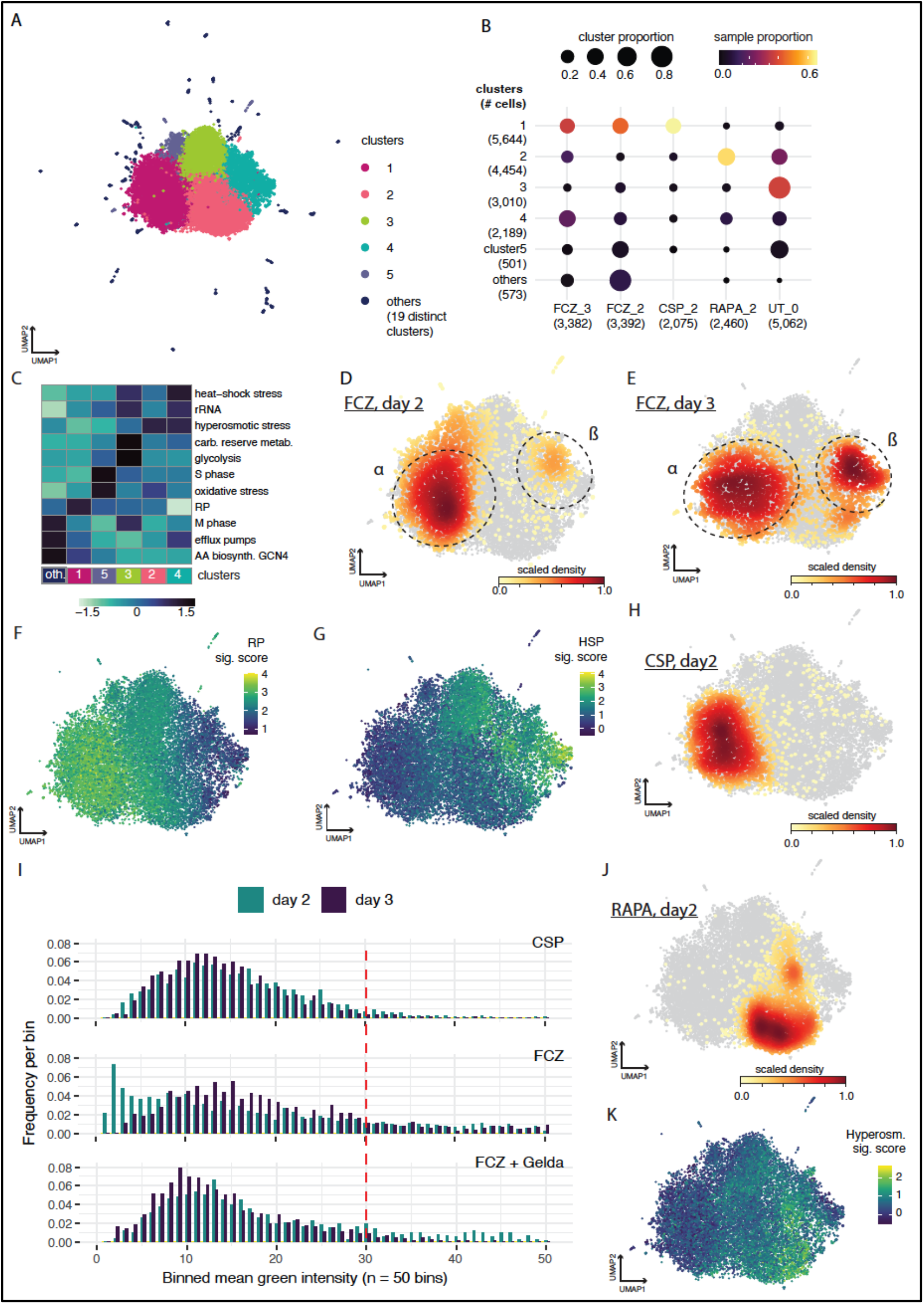
Cell profiles after challenges with different antifungal compounds. **(A)** UMAP embedding of untreated (UT) cells in addition to cells treated with fluconazole (FCZ) (days 2 and 3), rapamycin (RAPA) (day 2) and caspofungin (CSP) (day 2). Leiden clustering identified 5 main clusters and 19 sparsely populated “comets”, as color-coded. **(B)** Rows correspond to clusters and columns correspond to drug_day conditions. Dot diameter is proportional to the fraction of cells from each condition for a given cluster. The dot color is proportional to the fraction of cells from each cluster for a given condition. Numbers in parentheses indicate the total cell count in clusters and drug/timepoints conditions. **(C)** A heatmap depicting the level of activation (VISION z-scores) of different signatures, analogous in methodology to Figure 2B. **(D-J)** The UMAP embedding from (A). Color bar depicts: **(D,E,H,J)** density of cells for each condition **: (D)** FCZ at day 2, **(E)** FCZ at day3, **(H)** CSP at day 2 and **(J)** RAPA at day 2 ; or **(F,G,H)** signature scores for the **(F)** RP, **(G)** heat-shock stress and **(K)** hyperosmotic stress signatures. **(I)** Histograms depicting expression of HSP70 in CSP, FCZ and FCZ with geldamycin at days 2 (green) and day 3 (blue).

Genes overexpressed in the collection of 19 comets compared to the five major clusters include *MET10*, a bifunctional cysteine synthase/O-acetylhomoserine aminocarboxypropyl transferase which localizes to the cell wall, *GCN4* a transcriptional activator of the general amino-acid response in addition to *ILV2* and *ILV5* which encode for enzymes involved in branched-chain amino acid biosynthetic pathway (**Figure 3 – figure supplement 2A ; Method 9;** Bayes factor > 2.5 and proportion of non-zero value > 0.2). The ILV pathway is well studied as a fungal amino acid starvation therapeutic and *ILV5* is recognized as a component in the *C. albicans* amino-acid starvation response (Kingsbury and McCusker, 2010; Tripathi et al., 2002). It is regulated by the transcriptional activator *GCN4* in both *S. cerevisiae* and *C. albicans* (Hinnebusch, 2005; Natarajan et al., 2001; Tripathi et al., 2002). Signature analyses confirmed high expression of *GCN4* target genes encoding amino-acid biosynthetic enzymes as well as high expression of genes involved in drug efflux and low expression of ribosomal RNA (rRNA) in the 19 comet clusters (**Figure 3C**). Although the number of cells per comet cluster is small (from 7 to 126 cells), individual molecular marker genes could be identified in comets and some of these markers have been reported to be involved in FCZ tolerance or resistance (**Figure 3 – figure supplement 2B ; Methods 9;** top 5 with Bayes factor > 3 and proportion of non-zero value > 0.2). These included cell wall organization and adhesion (*ADA2, TPK2, LMO1, KRE9, RDH54, DFI1*) and, chromosomal and cytoplasmic division (*AXL2, HOS3, RDH54, RFA2 RIM15, NPL4, CDC5 BUB2*).

To further compare FCZ cells that are part of comets with FCZ cells located in the main clusters 1-5, we performed differential expression analysis between the pseudo-bulk for cells in comets and a pseudo-bulk cluster formed from the rest of FCZ cells in the main clusters (**Methods 9)**. This analysis identifies 60 genes which are almost all overexpressed in comets compared to other FCZ-treated cells (DESeq2, FDR < 0.1; **Figure 3 – table supplement 1A**). Gene ontology analysis points to an enrichment of genes involved in the regulation of DNA replication, actin cytoskeleton, plasma membrane organization and mitotic ring assembly, all highly expressed in the comet clusters (*NHP6A, HHF22, HHF1, SBA1, NCE102, KIN2, RVS167, CDC12*) (**Figure 3 – table supplement 1B, figure supplement 2C)**. This is consistent with the fact that FCZ impairs bud formation and is observed concomitantly with either DNA replication or duplication of the spindle pole body (Harrison et al., 2014). This effect can lead to the formation of trimeras and aneuploidy. Using the sc-data, we searched for evidence of aneuploidy by calculating the percentage of reads assigned to genes present in each chromosome. We observed that cells in cluster 16 have a significantly higher proportion of reads assigned to genes in chromosome 2, which could indicate trisomy or local chromosomal amplification (**Figure 3 – figure supplement 2D**).

### 5. Cells display distinct survival responses to fluconazole

Fluconazole (FCZ) targets *ERG11* in the sterol pathway, disrupting sphingolipid biosynthesis and membrane integrity (Odds et al., 2003; Thamban Chandrika et al., 2018; Wertheimer et al., 2016). The clinical relevance of FCZ in addition to findings that drug tolerance to FCZ evolves quickly and is clearly evident only three days post treatment (Gerstein and Berman, 2020) motivated us to profile *C. albicans* cell subpopulations at 2 and 3 days post-exposure. From the sc-transcriptional profiles, we observe a strong bifurcation in the surivor cell population already by day 2, suggesting that cells respond in at least two distinct ways to the FCZ challenge (**Figure 3D**). This bifurcation becomes more striking by day 3 post-treatment (**Figure 3E**). Almost every surviving FCZ-treated cell appears in either one of these two states. The first survival state lies at the “left” cluster 1-darkpink of **Figure 3A**, termed the α response in **Figure 3D**. This region is characterized by high RP expression and an absence of either heat-shock or hyperosmotic stress genes (**Figure 3C,F,G,K**). Signature analysis also suggests that the α group of survivors lowly express genes involved in the glycolytic and carbohydrate reserve metabolic pathways in addition to S-phase histones. However they have moderate to high expression of the GCN4-mediated response that activates amino acid biosynthesis (**Figure 3C – figure supplement 3A**).

The second survival state lies at the right of **Figure 3D,E**, termed the β response. These FCZ cells are characterized by a low expression of genes coding for RP and a high heat-shock stress response (**Figure 3C,F,G – figure supplement 3A**).

A comparison of pseudo-bulk profiles of FCZ α response cells (from cluster 1-darkpink) versus β response cells (from cluster 4-turquoise) highlights 797 differentially expressed genes (DESeq2, FDR < 0.1; **Methods 9**; **Figure 3 – table supplement 2A**). Some 230 of these genes are overexpressed in α cells with half of those involved in protein translation **(Figure 3 – table supplement 2B, figure supplement 3B)**. Many highly expressed genes within the α response are involved in rRNA processing and mitochondrial cellular respiration.

Cells in the β response overexpressed 567 genes involved in cell wall organization (e.g., *CRH11*, *BUD7*, *CHS5*), cell adhesion, morphology and virulence (e.g., filamentous growth, biofilm formation incl. *CPH2*) **(Figure 3 – table supplement 2, figure supplement 3B)**. Interestingly, β cells also highly express genes involved in the unfolded protein response (e.g., *HSP70, YHB1*) genes that promote drug tolerance (e.g., *HSP90, GZF3* (Delarze et al., 2020; Rosenberg et al., 2018), in addition to *CCH1* (Liu et al., 2015)*, HSP21* (Mayer et al., 2013)*, HSP70* (Nagao et al., 2012), and *RIM101*(Garnaud et al., 2018).

In summary, isogenic cells, which manage to survive treatment with FCZ, respond in one of two ways. The α response involves an upregulation of protein translation, rRNA processing and mitochondrial cellular respiration whereas the β response involves processes that assist the cell to cope with the effect of FCZ in addition to several stress responses.

### 6. Subinhibitory dose of caspofungin triggers the α response

We next investigated whether the bifurcation observed with FCZ is also present with a second antifungal CSP, an echinocandin used as a first-line clinical treatment. CSP acts by inhibiting β-1,3 glucan synthesis and disrupting the fungal cell wall (McCormack and Perry, 2005). Although it is effective against most *Candida* species, one well-known mechanism of echinocandin resistance involves point mutations in “hot spot” regions of FKS-encoded subunits of a glucan synthase (Perlin, 2015; Pristov and Ghannoum, 2019). Tolerance to CSP increases with larger amounts of cell wall chitin, which could be the result of aneuploidy and rearrangement of chromosome 5 (Yang et al., 2017). Although fungicidal CSP has a different mechanism of action as fungistatic FCZ, similar mechanisms of tolerance were found linked to Hsp90, the calcineurin pathway (Singh et al., 2009) and the pH-responsive RIM pathway (Garnaud et al., 2018).

We treated a *C. albicans* population with a subinhibitory concentration of CSP, well below its MIC_50_ (1 ng/mL) because higher doses of CSP triggered significant cell aggregation which made the cells unsuited for sc-profiling. At day 2, the vast majority of cells localized to cluster 1- darkpink, where α FCZ cells also reside (86 % of all CSP, **Figure 3B,H**).

We asked if there were differences between CSP and FCZ cells in the α state. This analysis identified only one gene significantly differentially expressed (FDR < 0.1) suggesting that the α response is overall very similar for both anti-fungal treatments, at least at the chosen dosage levels and day 2 time point.

Unlike FCZ, there was no evidence of a bifurcation for CSP at day 2. This could be due in part to our choice of a low dosage level, a parameter that has been previously shown critical determining the cellular response to CSP in *C. albicans* (Enjalbert et al., 2006, 2003). There is nevertheless some evidence of heterogeneity in cellular response to CSP with approximately 6% (N=136) of CSP-treated cells found in either the RAPA-enriched cluster 2-pink (11%), UT-enriched cluster 3-green (4%) or in HSP-stress cluster 4-turquoise (3%) (**Figure 3C**).

It is possible that higher concentrations of CSP in addition to longer time courses could induce a bifurcation analogous to FCZ with emergence of β cells characterized by high expression of heat-shock protein coding genes. Since our analysis above highlights *HSP70* as a strong marker of the β response, we analyzed microscopy images of our GFP-tagged *HSP70* cells exposed to CSP or FCZ across two and three days. While a significant fraction of cells treated with FCZ highly express Hsp70 at day 2 and 3 confirming the presence of β cells, no similar trend was observed in cells treated with CSP at either days (**Figure 3I**). We further analyzed microscopy images of cells treated with both FCZ and the Hsp90-specific inhibitor Geldamycin; although some cells still exhibit high expression of Hsp70 at day 2, Hsp70 expression was significantly reduced in most cells at day 3 (**Figure 3I**).

### 7. Rapamycin survivors exhibit the highest expression of hyperosmotic stress and lowest expression of ribosomal proteins

The TOR (target of rapamycin) pathway controls many processes including protein translation, autophagy, apoptosis and cell growth in response to nutrient availability (Bastidas et al., 2009; Cruz et al., 2001; Heitman et al., 1991; Schmelzle and Hall, 2000). Inactivation of TOR by RAPA represses the expression of RP (Powers and Walter, 1999), triggers various cellular mechanisms aimed at overcoming nutrient stress (Cruz et al., 2001), and disrupts sphingolipid biosynthesis which in turn can induce cell wall or plasma membrane stress (Teixeira and Costa, 2016). Although *C. albicans* is markedly sensitive to RAPA (Baker et al., 1978), it has weak antifungal properties when used in isolation (Baker et al., 1978; Tong et al., 2021). However, RAPA treatment concomitantly with azoles including FCZ has a synergistic effect (Tong et al., 2021) resulting in the ablation of drug tolerance (Rosenberg et al., 2018). For this reason, we sc-profiled *C. albicans* populations challenged with a concentration of RAPA which does not visibly impact growth (0.5 ng/ml; Figure 3 – figure supplement 1A) but we know enables inhibition of FCZ tolerance (Rosenberg et al., 2018).

Cells treated with RAPA localize primarily to cluster 2-pink and to a lesser extent to cluster 4- turquoise in the β state (**Figure 3B, J**). We previously observed that β FCZ cells have high activation of genes in the heat-shock stress response. This is also observed to be true for RAPA treated cells, but there is also a very high hyperosmotic stress response (**Figure 3K**). In fact the elevated hyperosmotic stress response is found in all RAPA cells regardless of their cluster classification (**Figure 3 – figure supplement 3C**, light blue curves). In particular, RAPA cells have high expression of the sodium pump *ENA2*, and the glycerol-3-phosphate dehydrogenase *GPD2* (**Figure 3 – figure supplement 4A**). Both genes are orthologues of hyperosmotic responsive genes in *S. cerevisiae* and are induced in *C. albicans* in response to hyperosmotic stress (Enjalbert et al., 2003), suggesting that activation of genes involved in hyperosmotic stress is very specific to RAPA treatment.

Finally, although the RAPA-enriched regions appear to have moderate to low RP (**Figure 3F,J**), careful analysis revealed that RAPA cells have the lowest RP expression compared to cells from any other conditions (**Figure 3 – figure supplement 4B**). This is expected since RAPA inhibition of TOR signaling is known to decrease expression of RP genes (Powers and Walter, 1999; Schmelzle and Hall, 2000). As expected, no RAPA cells were therefore found in the α state characterized by high RP.

### 8. FCZ survivors exhibit increased drug tolerance already by day 2 which stabilizes at day 3

We asked if the degree of drug tolerance and resistance exhibited by FCZ survivors changes across the time series extended to six days. Via a series of disk diffusion assays with FCZ detailed in **Appendix 1**, we measured changes in tolerance and resistance from 1 to 6 days post-exposure. A statistically significant increase in drug tolerance is observed between days 1 and 2 (p < 0.001, Kruskal-Wallis χ^2^ test). Drug tolerance again rises from days 2-3. Then, the level of drug tolerance remains the same from days 3 to 6. There was a small but statistically significant increase in drug resistance at day 2 post-treatment. Overall, these results confirm that survivors are enriched for cells capable of mounting a successful tolerance response, and that the vast majority of these cells have not developed long-term drug resistance.

### 9. Revisiting FCZ survivors at day 6

Given the dynamics observed for FCZ survivors between day 2 and day 3, we asked how these subpopulations would continue to change up to 6 days post-treatment. To investigate this, FCZ day 3 survivors were resuspended in YPD and sc- profiled three days later (FCZ day 6, **Figure 1A, Methods 1**). **Figure 4A** depicts the UMAP embedding of the sc-transcriptional profiles restricted to the FCZ-3 and FCZ-6 survivor populations. Leiden unsupervised clustering identifies four main clusters with some comets as similar to those described above. However, only five cells from day 6 are found in comets, we excluded comets in the downstream visualizations. Using **Figure 4B,C,D** allowed us to relocate the day 3 α response, which is mostly found in cluster 2-salmon with high RP expression, and the β response, which is captured by cluster 4-kaki with high expression of the heat-shock stress signature.

**Figure 4.**
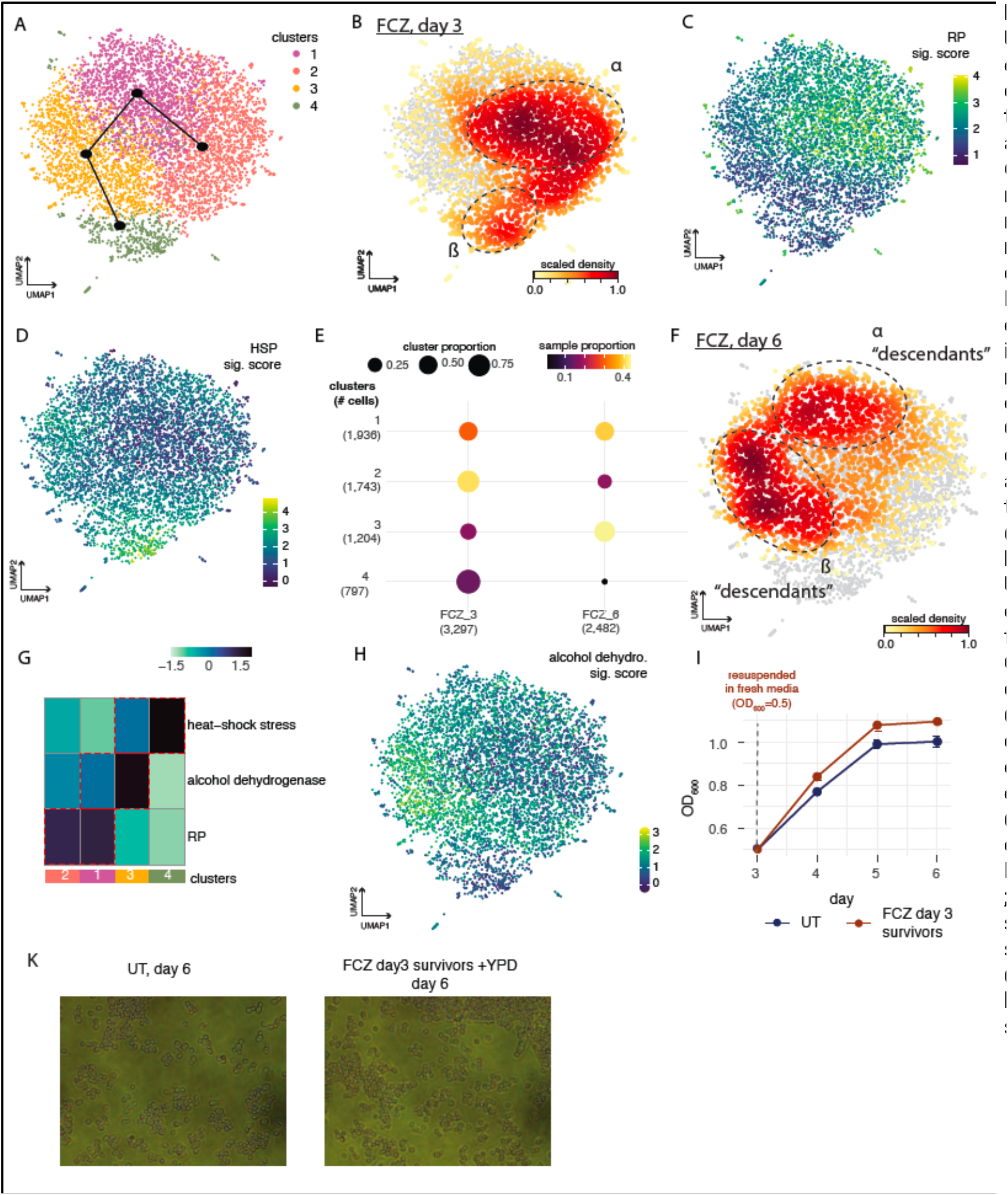
**(A)** UMAP embedding of FCZ- treated cells at day 3 and 6 (after resuspensio n in fresh media at day 3). Leiden clustering identified 4 main clusters. Clusters are ordered along a trajectory (black). **(B- D, F, H)** The UMAP embedding from (A). Color bar depicts: **(B,F)** density of cells for each condition **: (B)** FCZ at day 3, **(F)** FCZ at day 6; or **(C,D,H)** signature score for the **(C)** RP, **(D)** heat-shock stress, and **(H)** alcohol-dehydrogenase gene signatures. **(E)** Rows correspond to clusters and columns correspond to drug_day conditions. The diameter of the dot is proportional to the fraction of cells from each condition for a given cluster. The color of the dot is proportional to the fraction of cells from each cluster for a given condition. **(G)** A heatmap depicting the average level of activation (VISION z-scores, color bar) of different signatures across clusters presented in (A), analogous in methodology to Figure 2C.**(I)** Growth curves for untreated (UT) and FCZ day 3 survivor resuspended in fresh media as measured by OC_600_ across days. **(K)** Microscopy images at day6 of UT cells (left) and FCZ day3 survivors (right) resuspended in YPD.

Approximately 70% of day 6 survivors have shifted to clusters 1-purple and 3-yellow (**Figure 4E,F**). Cluster 3-yellow has strong expression of heat-shock stress and low expression of RP genes (**Figure 4G**), consistent with the β cells at day 3. This may indicate that cluster 3-yellow survivors at day 6 are the “descendants” of day-3 β cells. The remaining day 6 survivors are almost exclusively found in cluster 1-purple, which has high expression of RP and no expression of the heat-shock signature. This pattern is characteristic of the α cells at day 3 and similarly may indicate that cluster 1-purple survivors at day 6 might be the “descendants” of day 3 α cells. These hypotheses were supported by the lineage structure inferred by computational trajectory analysis (**Figure 4A**, black tree; **Methods 8**, slingshot analysis). Overall, these results indicate that “descendants” originating from both α and β populations survive to day 6.

The significant difference in the relative size of the α and β populations at day 3 disappears by day 6. This may suggest that β cells are more fit to proliferate, even if both the α and β responses represent viable long-term survival options. We also observe that cells derived from the β response have the highest expression of alcohol dehydrogenase enzyme-coding genes at day 6 (**Figure 4H**). This process is involved in ethanol utilization under growth conditions. After resuspension of day 3 FCZ survivors in fresh media, OD600 measurements confirmed that FCZ day3 survivors resuspended in fresh media are not only fit to proliferate, but do so at a rate faster than UT cells (**Figure 4I**).

Pseudo-bulk analysis identified 868 genes differentially expressed between the day 6 α and β “descendants” (FDR < 0.1; **Figure 4 – table supplement 1A**). A very high number (N=315) of these genes were also previously identified as differential between the α and β cell populations at day 3 (Fisher’s exact test p < 0.0001). This provides additional evidence that the α and β cell populations at day 6 likely originate from α and β cell populations at day 3. Not surprisingly then, many biological processes characterizing the α and β descendants at day 6 are also enriched in genes overexpressed in α or β cells at day 3 (**Figure 4 – table supplement 1A,B**).

For both day 3 and 6 survivors, genes overexpressed in α cells are involved in translation processes and rRNA processing (**Figure 5,** center **– table supplement 1A**). Within these rather broad GO categories, we can identify several specialized biological processes only enriched at day3 or only at day 6 (**Figure 5**, left and right column, respectively). In particular, GO enrichments for day-6 α descendants include regulation of translational termination and elongation (GO group 1), reinitiation of translation (GO group 2,3), tRNA and rRNA catabolic processes (GO group6), nucleolytic cleavage of rRNA to generate mature rRNA (GO groups 8, 9), and ribosomal subunit export, biogenesis and assembly (GO group 10-12). In contrast, specific processes enriched in α cells at day 3 are related to mitochondria cell respiration (GO groups 4,5).

**Figure 5.**
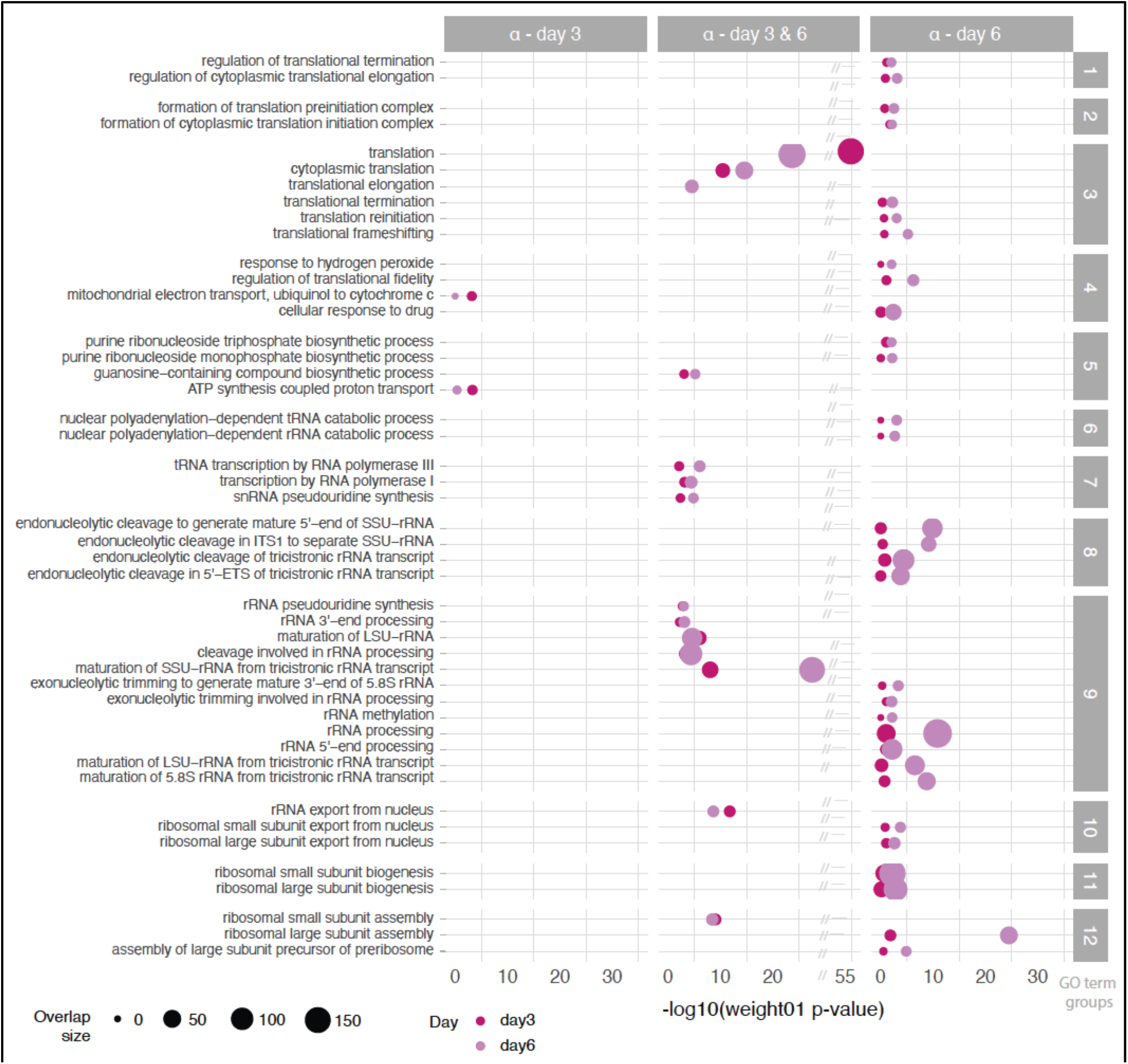
GO enrichments for genes overexpressed in FCZ α cells compared to FCZ β cells at day 3 and day 6 (color) (N = 230 and 474 genes, respectively ; FDR <0.1). GO terms are grouped into clusters (rows) based on semantic similarity and into columns depending if the term was significant for the day 3 gene list only (left), the day 6 gene list only (right) or for the gene lists at both days (center) (weight01 fisher p-value < 0.01). The size of the dot is proportional to the number of genes in the list which overlap with the corresponding GO term.

Similarly, genes overexpressed in β cells at both day 3 and 6 are enriched for several processes involved in metabolic and nutrient-limited condition response (e.g., carbohydrate transport, piecemeal microautophagy of the nucleus, acetate catabolism, gluconeogenesis; GO groups 1,2,12,16; **Figure 6**) while a few metabolic processes related to glucose transport, glycolytic fermentation and amino-acid are enriched only at day 6 (**Figure 6**). Although heat-shock protein-coding genes (*HSP70, HSP78, HSP104*) are overexpressed at both days, we only observe significant activation of the cellular response to unfolded/misfolded protein and to heat at day 3 (GO groups 4, 8, **Figure 6**). Other processes related to filamentous growth (GO groups 5, 6) and biofilm formation (GO group 5) are enriched in β cells only at day3 (**Figure 6**).

**Figure 6.**
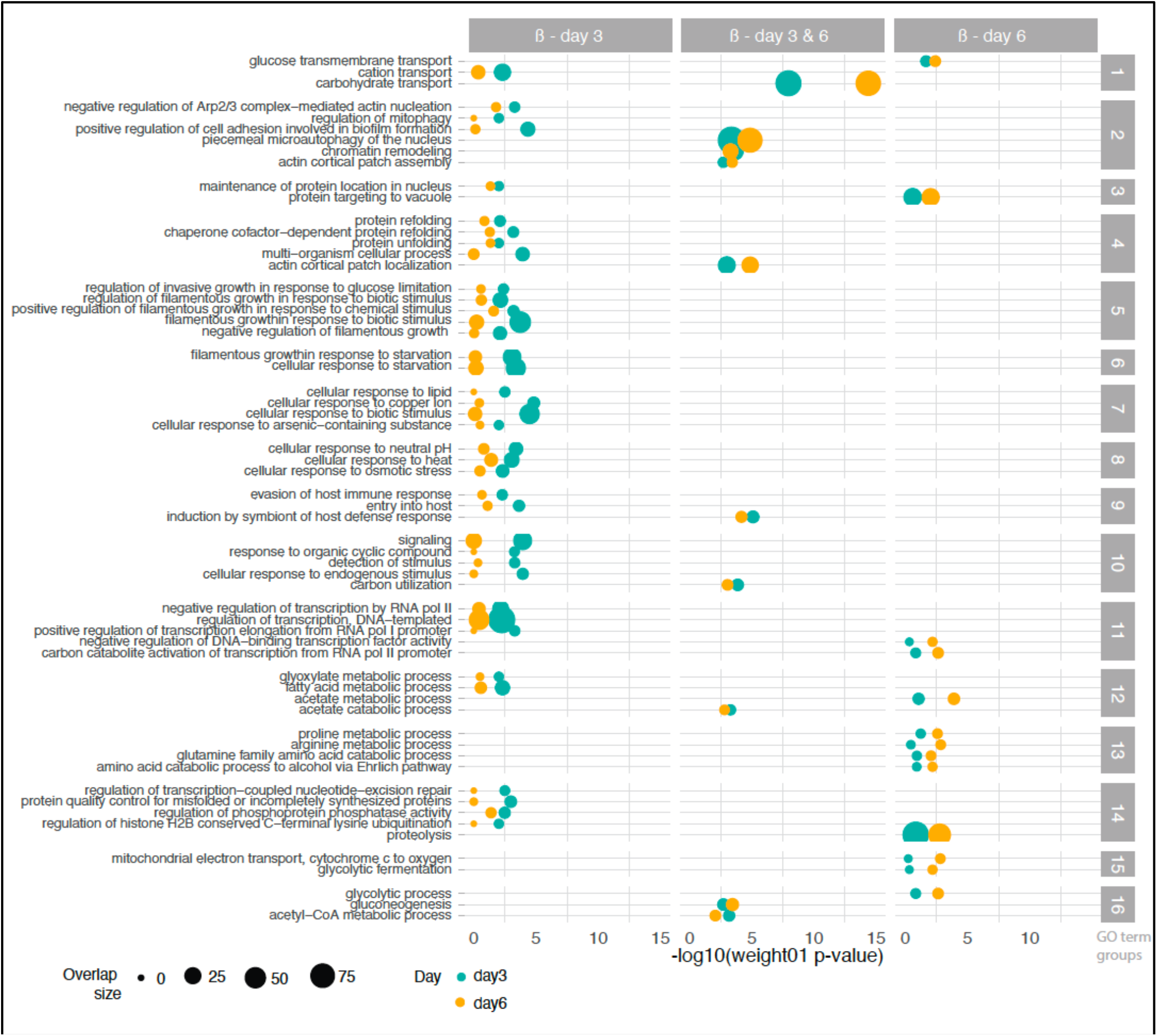
GO enrichments for genes overexpressed in FCZ β cells compared to FCZ α cells at day 3 and day 6 (color) (N = 567 and 394 genes, respectively ; FDR <0.1). GO terms are grouped into clusters (rows) based on semantic similarity and into columns depending if the term was significant for the day 3 gene list only (left), the day 6 gene list only (right) or for the gene lists at both days (center) (weight01 fisher p-value < 0.01). The size of the dot is proportional to the number of genes in the list which overlap with the corresponding GO term.

Overall, these results indicate that both populations are viable survival routes. The observation that FCZ-treated cells exhibit higher growth than UT cells is likely explained by the fact that β cells express metabolic and cellular processes related to cell growth and expand quickly in the presence of fresh media. Although many processes that are differentially expressed between the α and β populations are common to both timepoints, it is the day 6 α descendants which exhibit reinitiation of translation. Likewise, it is the day 6 β descendants that express growth-related processes likely enabled by the expression of tolerance response programs.

## Discussion

Single cell (sc) approaches provide capacity to explore aspects of phenotypic heterogeneity in microbial populations in several important dimensions. For example, using unsupervised analysis techniques with profiles of thousands of individual cells, we are able to detect subsets of individuals that share common molecular features but which are distinct from the remaining cells. Although the number of transcripts identified per cell in our *C. albicans* setting is far fewer than previous nanoliter droplet-based single cell profiling studies of mammalian systems, dimension reduction techniques (i.e., a latent space of a variational autoencoder) and subsequent clustering were still able to identify three subpopulations of untreated (UT) prototrophic SC5314 *C. albicans* cells.

The sc-transcriptomic profiles established that each of the three UT subpopulations exhibited significant heterogeneity in expression of genes involved in cell cycle phase in addition to metabolic and stress responses. Specifically, we observed that glycolysis and several heat-shock genes are upregulated during the M phase, oxidative stress genes are upregulated in a distinct subset of cells likely at the early stage of cell cycle, and ribosomal proteins (RP) are upregulated during the S phase (captured by the simple model depicted in **Figure 2D**). Many of these associations are well supported by the literature (Chiu et al., 2011; Finkel and Hwang, 2009; Hossain et al., 2021; Senn et al., 2012), and together provide evidence of the efficacy of our sc- assay to detect cell-to-cell heterogeneity even within untreated *C. albicans* cultures.

To further validate the existence of the subpopulations, we developed dual fluorescence reporter strains that tagged Hsp70 with GFP and Ttr1 with RFP, and grew cells under the same conditions as per the sc-profiling assay. We again observed a significant mutual exclusivity where cells expressed either Hsp70 or Ttr1, but not both at high levels. Microscopy analysis also established that the vast majority of all cells (∼98%) were observed to be in the yeast white morphology, negating the possibility that a morphological switch is responsible for the distinct subpopulations. DNA-level sc-profiling has not been established yet for microbial systems, however bulk DNA profiling of the SC5314 strain identified the same number of single nucleotide polymorphisms and structural variants as a previous effort from Cuomo and colleagues (Cuomo et al., 2019). The degree of penetrance of any single mutation or copy number variant is too small to explain the size of the observed clusters within the timeframe of our experiments, suggesting that these distinct subpopulations are very unlikely due to pre-existing genomic differences between individuals. We note however that although the cell cycle and their associated signatures including stress and metabolic responses are the main drivers of variability in UT cells, the magnitude of these transcriptional signatures is much lower than later analyses where cells show more intense transcriptional perturbations in response to antifungal challenge.

After treatment with FCZ, we observed more than a dozen small clusters of cells “radiating out” from the center of the UMAP. Differential expression analysis identified genes involved in amino acid biosynthesis, efflux activity and chromosome segregation and mitosis as overexpressed in many of these “comets”. Although the mechanism remains unknown, exposure of *C. albicans* to FCZ was previously associated with decreased pool sizes of intermediates of amino-acid biosynthesis (Katragkou et al., 2016). The altered efflux of the cell membrane may have increased the release of amino-acid from the cells, as previously observed in auxotrophs (Yu et al., 2022) or the amino-acid starvation response could reflect the alteration of the translational machinery (rRNA, RP) when cells are under extreme stress (Hinnebusch, 1994; Tournu et al., 2005). Overall, “comets” may represent the emergence of trimeras with unstable genomes previously identified after FCZ treatment (Harrison et al., 2014) which might have arisen as a response to amino-acid starvation .

The application of unsupervised dimension reduction/clustering to sc-profiles of FCZ- challenged *C. albicans* identified distinct subpopulations across six days post-treatment. In particular, we observed a bifurcation at day 2 into the so-called α and β responses. At day 2, the majority of FCZ treated cells are in the α cluster, but within 24 hours the two clusters are approximately balanced. By day 6 the β cluster appears to be more prevalent. Microscopy again confirmed that the vast majority of cells remained in the white morphology. We performed bulk DNA-level profiling of these populations and observed massive and near genome-wide instability with multiple loss, loss of heterozygosity and amplification events scattered throughout the genome (**Figure 2 - supplement figure 5**). This instability appears to peak at day 3 and falls to near normal levels by day 6 . Attempts to assign genomic aberrations to either the α and β response were inconclusive, suggesting perhaps that both responses contain unstable cells at these early time points. We observed only a small increase in drug resistance for these FCZ populations at day 6 compared to control. However, there was a large increase in drug tolerance between days 1 and 2 that persists to day 6. This suggests that survivor populations have not evolved resistance *en masse* by day 6, however there has been some selection for cells that are capable of mounting an efficacious tolerance response.

The α response, which is more prevalent at day 2, is characterized by high expression of ribosomal protein (RP) and ribosomal RNA (rRNA) processing genes. Ribosomal biogenesis requires a balance in the synthesis of RP and rRNA, and disequilibrium leads to proteotoxic stress and RP aggregation in yeast (Tye et al., 2019). In turn, this stress and aggregation induce the so-called Ribosome Assembly Stress Response (RASTR) in yeast which involves the heat-shock transcription factor Hsf1. Hsf1 in turn upregulates *HSP90* and other genes involved in protein folding including *HSP70* (Albert et al., 2019). The transcription factor Ifh1 is also central in RASTR. It is highly sensitive to RP aggregation which causes it to dissociate from RP genes. This in turn has the effect of suppressing RP expression thus providing protection against toxic protein build-up and restores the ability of *S. cerevisiae* cells to grow (Albert et al., 2019).

*C. albicans* has a highly conserved orthologue of *HSF1* from *S. cerevisiae* (official name *CTA8*, with alias *HSF1*). In addition to *HSF1*, the proteostasis networks and ribosome assembly are also highly conserved, indicating that the conditions which disrupt ribosome assembly in *S. cerevisiae* may also drive proteostasis collapse in *C. albicans* (Tye et al., 2019). In our data, *HSP70,* and *HSP90* as well as the disaggregases *HSP21*, *60*, *78*, and *104* are more than 2-fold over-expressed in the β response compared to the α response. Pathway analysis of the differentially expressed genes also highlights a statistically significant enrichment of unfolded/misfolded protein-related gene ontology terms. In addition, the β response has low RP expression and no enrichment for rRNA-related processes. This suggests a model whereby cells, when exposed to FCZ, initially transit into the α response characterized by imbalances in the synthesis of RP and rRNAs. This triggers the RASTR response which shuts down RP and restores the ability of cells to grow. The molecular profile of the β cells is consistent with this transition: in addition to the low expression of RP genes and rRNAs, β cells highly express the RASTR induced genes including *HSP90* (Rosenberg et al., 2018) and *HSP70* (Nagao et al., 2012) in addition to many other drug tolerance-related genes including PCK1 (Wuyts et al., 2018), *GZF3* (Delarze et al., 2020; Rosenberg et al., 2018)*, CCH1* (Liu et al., 2015)*, HSP21* (Mayer et al., 2013), and *RIM101* (Garnaud et al., 2018). Moreover, this transition from the α to β state is consistent with the change in the relative population size these two subpopulations just 24 hours later. Together this suggests that the β cells likely represent the subpopulation of *C. albicans* which have mounted a successful drug tolerance response.

Although cells in both α and β states survive to day 6 after resuspension in fresh media, β cells are more prevalent. This is consistent with the higher proliferative capacity of the β cells, since β cells exhibit increased expression of an alcohol dehydrogenase response, suggesting that the surviving β cell population is expanding. The unfolded protein response is highly expressed only at day 3, although a subset of HSP-related genes remain over-expressed until day 6. This suggests that at day 3 the RASTR rescue has succeeded and the cells, when resuspended in fresh media, have now mounted a successful drug tolerance response and are proliferating. At day 3, β cells also highly express genes involved in filamentous growth, adhesion and biofilm formation which is consistent with a potential role of Hsf1 (Hahn et al., 2004; Leach et al., 2016, 2012; Veri et al., 2018). Lastly, we highlight that our data suggests that at least some α cells are able to survive without this RASTR-mediated transition to the β response. Although this state might be less favorable than the β response, we do observe a significant proportion of descendants from α cells that reinitiate translational machinery after resuspension in fresh media.

Cells treated with rapamycin (RAPA) were depleted from the α response characterized by high RP expression, and co-clustered closely to β response cells. This is perhaps not surprising given that by targeting Tor1, RAPA shuts down ribosomal biogenesis. Concomitant exposure to RAPA and FCZ is known to ablate drug tolerance (but not drug resistance) (Rosenberg et al., 2018). Our results suggest that this might occur through the inhibition of the α response so cells cannot transitioned to the tolerant β state. Of note, RAPA specifically triggered the expression of genes involved in hyperosmotic stress. Previous work established that the Tor1 pathway controls phase separation and the biophysical properties of the cytoplasm by tuning macromolecular crowding (Delarue et al., 2018). Hyperosmotic stress response to RAPA exposure might reflect disruption of the biophysical properties of the cytoplasm, altering the osmotic balance in cells.

Finally, cells treated with caspofungin (CSP) co-clustered very closely with FCZ α response cells, although at day 3 there was no evidence of CSP cells in the β response. The similarity in the early response is perhaps surprising given that CSP and FCZ have different mechanisms of action (i.e., disrupting cell wall biosynthesis versus disrupting ergosterol biosynthesis). It will be interesting to determine the universality of the α response in future antifungal profiling efforts and the potential transition to the proliferative tolerant β state. Together this supports the notion that drug tolerance exploits stress response strategies that are enabling survival despite the continued ability of the drug to interact with its target.

## Conclusions

This is the first single cell study of *C. albicans* populations with a nanoliter droplet-based assay modified for the fungal setting. The assay was relatively inexpensive with few difficulties or failed runs, however optimizations to improve the capture rate would likely allow us to peer more deeply into the community structure, and would simplify the downstream data science analyses. These analyses relied on gene signatures from the literature that were often derived from previous bulk transcriptome studies. Studies like ours help to refine these signatures by highlighting when specific genes, pathways and responses are ubiquitously expressed by all cells versus when they are expressed by only a subclonal population. The ability to study cellular trajectories across these populations in a high-throughput manner also provides new perspectives on time-sensitive processes such as the emergence of drug tolerance and resistance. Targeting β cells, or genes and pathways that support the α to β transition are interesting avenues to block drug tolerance. If the α response generalizes to other drug treatments, interrupting the α to β tolerance transition could represent a novel therapeutic direction across different anti-fungal treatments.

## Materials and Methods

### 1. Strains, media, and drug treatment

#### a. Colonies for sc-RNA- bulk RNA- and DNA-sequencing

*C. albicans* SC5314 cells were streaked out from glycerol stocks in -80°C on YPD agar plates (2% D-glucose, 2% peptone, 1% yeast extract, 0.01% uridine, agar 2%) and incubated at 30°C for 48 hours. Afterwards, a single colony of cells was transferred into YPD liquid media (2% D-glucose, 2% peptone, 1% yeast extract, 0.01% uridine) and incubated at 30°C for 12-16 hours.

#### b. Further preparation for sc-profiling via DROP-seq

An aliquot of 10^8^ cells was taken and served as our colony. For untreated cells, we introduced 1ml of RNAlater (Sigma # R0901) and froze the resultant colony at -20°C for later use in the DROP-seq. For treated cells, we performed the following approach to ensure a sufficient number of mid-log phase cells for profiling with different drugs and multiple time-points: cells were pelleted and resuspended in 1ml of YPD. Then, 250μl of this suspension was combined with 15ml of fresh YPD and placed in a shaker incubator at 30°C until they reached an OD600 of ∼0.6. Finally, on the order of 10^8^ of these cells were placed in 10ml of YPD. Each suspension was then subjected to drug treatment (**Figure 1 – figure supplement 1**).

A concentration of 1 µg/mL was chosen for fluconazole (FCZ, Sigma #F8929) representing a moderate dosage relative to their reported MIC50 levels (Diaz et al., 2016; Girmenia et al., 2000; Pfaller et al., 2006; Shokohi et al., 2016; Tong et al., 2021). A concentration of 1ng/ml was used for caspofungin (CSP, Sigma #SML0425), a compound that interrupts cell way biosynthesis (Cappelletty and Eiselstein-McKitrick, 2007; Stevens et al., 2004; Yang et al., 2017). This is well below its reported MIC50 levels and was chosen in order to ensure a sufficient number of non-aggregated survivors to generate single cell profiles. A subinhibitory concentration of Rapamycin (RAPA) was chosen (0.5 ng/ml) based on previous studies that establish such levels are sufficient to generate a fungicidal synergistic effect when given concomitantly with FCZ (Rosenberg et al., 2018). Each drug was delivered to the individual colonies and incubated at 30°C for 48 or 72 hours. We also generated sc-transcriptional profiles of FCZ survivors after six days. For this population, day 3 survivors were washed twice, and resuspended at a concentration of ∼10^7^ in 1ml in fresh YPD.

At each timepoint, colonies were collected and strained (pluriStrainer® 20 µm, pluriSelect) before placement in fresh tubes. Straining was done in order to minimize the likelihood of clogging in the microfluidic due to rare but large hyphae and pseudohyphae morphologies. We observed that germ tubes up to four times the length of the mother cell can still be processed for DROP-seq analysis. Such cells are well within the hyphal transcriptional profile (Nantel et al., 2002), suggesting that our results may contain some profiles of hyphae and pseudohyphae cells. After filtering, the vast majority of cells were in the yeast white morphology with less than 0.2% of cells were in a filamentous morphology (hyphae or pseudohyphae) after manual counting ∼100 microscopy images with an average of ∼50 cells per slide for each such population. All cultures yielded a sufficient population of survivors for downstream sc-transcriptional profiling, bulk transcriptional profiling, bulk DNA genomic profiling and/or microscopy. Colonies were washed with 1ml of RNAlater twice. Cells were then resuspended in 0.5 ml RNAlater and incubated at room temperature for 30 minutes before storage at -20°C until sc-profiling with DROP-seq.

#### c. Colonies for OD600 analyses

*C.albicans* SC5314 cells were grown in 5ml YPD liquid for 12 to 16 hours in a 30°C shaker incubator. We grew three biological replicates for each experiment. The next day, cells were diluted to OD600 0.1 in 50 ml YPD liquid and incubated again until they reached an OD_600_ of 0.5. Afterwards, two technical replicates were taken for each experiment. Cells were either untreated or treated with FCZ, CSP or RAPA with the same concentrations that were used for the DROP-seq experiments. The aliquots were placed in a 30°C shaker incubator and the OD600 was measured after 1, 2 and 3 days. On day 3, cells were washed with YPD twice and diluted to an OD600 of 0.5. Cells were placed in the shaker incubator and their OD600 were measured each day over the next three days.

### 2. Spheroplasts

The *C. albicans* setting required an optimized protocol for the removal of the cell wall and to induce stable spheroplasts for single-cell profiling. Towards this end, we experimented with different concentrations of zymolyase (0.1, 0.2 and 0.4U zymolyase (BioShop # ZYM002) with 10^7^ cells in 100 ul of sorbitol 1M) at different time points (incubated at 37 °C for 10, 20, 30 mins) before processing with the DROP-seq. To compare colonies grown under different conditions, cells were stained with calcofluor white and imaged using a Leica DM6000 microscope. We concluded that concentrations in the range 0.15-0.25U after 25 minutes are able to induce spheroplasts that remain sufficiently stable for processing with our DROP-seq.

### 3. Cell preparation for sc-profiling

At the time of DROP-seq profiling, an aliquot of 10^7^ (OD=0.68 in 660 nm) cells was separated from the colony in **Methods 1**, and washed three times with sorbitol 1M solution. The cells were then resuspended in 100μl sorbitol 1M + 0.25 U Zymolyase and incubated at 37 °C for 25 minutes (as per our findings in **Methods 2**). Next, the cells were pelleted and resuspended again in 0.5ml of cold and fresh RNAlater for five minutes. Now, the cells were washed (centrifuged and pelleted) with 1ml of washing buffer (1M sorbitol, 10mM TRIS pH 8, 100ug/ml BSA) three times. Finally, 10^6^ cells (OD=0.08 in 660nm) were resuspended in 1.2 ml of the washing buffer. This cell suspension was then used as input to the DROP-seq device.

Cell preparation generally follows the protocol given by Macosko et al. (Macosko et al., 2015) with some exceptions. Whereas Macosko et al. recommends a ratio of 100K mammalian cells to 120K beads for DROP-seq, we found that a ratio of 1M cells for 120K beads generated a sufficient yield of cDNA as per the Agilent Tapestation. Jackson et al. (Jackson et al., 2020) use 5M cells as input to the Chromium (10X Inc.) system. Furthermore, whereas Macosko et al. use 1ml of lysis buffer, we use 1.2 ml. Instead of 13 PCR cycles, we use 17 (Jackson et al. uses 10 cycles). Samples were sequenced using a NEXT-seq 500 (Illumina Inc.) following the standard Macosko et al. protocol set to yield an estimated 200 million reads/sample.

### 4. Quality control, basic processing and normalization of the single cell profiles

In general, all computations were performed using `Python` version 3.9.6 or the `R` version 4.0.4. Gene abundances were estimated from raw sequencing data using the end-to-end pipeline `alevin-fry` (He et al., 2021) which performs UMI deduplication and reduces the number of discarded (multimapped) reads. The pipeline utilizes a reference index covering the spliced transcriptome extracted from the latest version of *C. albicans* strain SC5314_A22 (haplotype A, version 22; GCF_000182965.3). The Unspliced-Spliced-Ambiguous (USA) mode was used to separately keep track of the types of transcripts from which UMIs are sampled. A gene-by-cell matrix for each sample was obtained by summing reads labeled as either “spliced” or “ambiguous” by `alevin-fry`.

We started by filtering genes from downstream analyses with a zero sum count (the count across all cells), as were cells with less than five, or more than 2000 transcript counts (**Figure 1 - figure supplement 1A**). Then, the `EmptyDrops` approach was used to exclude cells that were indistinguishable from background ambient RNA levels (Lun et al., 2018). `SCANPY` (Wolf et al., 2018), a python-based toolkit, and `SingleCellExperiment` (Amezquita et al., 2019), a `R/Bioconducto`r package, were used for data quality control, filtering of genes and downstream visualization. We removed genes (n = 869) that were expressed in less than 20 cells (**Figure 1 - figure supplement 1C**).

We then use `scVI` (Gayoso et al., 2021; Lopez et al., 2018) version 0.12.2, a Bayesian deep neural network architecture which implements a probabilistic model of mRNA capture and uses a variational autoencoder to estimate priors across batches and conditions. Models were built using default parameters for different grouping of samples : i) UT cells, ii) UT cells and CSP, RAPA, FCZ treated at day 2 (and 3), and iii) FCZ-treated cells at day 3 and 6. All models were adjusted for batch and library size. We trained `scVI`’s variational auto-encoder and stored the latent representation for visualization and downstream analyses. We reduced the inferred latent spaces to 2-dimensions via UMAP using the implementation of `umap-learn` (McInnes et al., 2020) in `SCANPY` (Wolf et al., 2018)(min_dist=0.3).

### 5. Bulk transcriptomics

Total RNA was extracted from FCZ treated cells at day 2 post-exposure, which were grown according to **Methods 1b**, using the Qiagen RNeasy mini kit protocol. RNA quality and quantity were determined using a Bioanalyzer (Agilent Inc.). Paired-end read sequencing (2 x 50bp) was carried out on a NextSeq500 sequencer (0.5 Flowcell High Output; Illumina Inc.). Raw reads were pre-processed with the sequence-grooming tool `cutadapt` (Martin, 2011) version 0.4.1 with quality trimming and filtering parameters: `--phred33 --length 36 --2colour 20 -- stringency 1 -e 0.1`. Each read pair was mapped against *C. albicans* strain SC5314_A22 (haplotype A, version 22; GCF_000182965.3) downloaded from the NCBI using `STAR` (Dobin et al., 2013) version 2.7.9a with the following filtering parameters: `--outSAMmultNmax 1 -- outSAMunmapped Within --outSAMstrandField intronMotif`. We obtained ∼13 millions reads of which 88% were uniquely mapped along the genome, The read alignments and *C. albicans* genome annotation strain SC5314_A22 (haplotype A, version 22; GCF_000182965.3) were provided as input to `featureCounts()` from the `Rsubread` package (Liao et al., 2019) version 2.4.3 to estimate gene abundances. The following parameters were used: `isPairedEnd=TRUE, countReadPairs=TRUE, requireBothEndsMapped=TRUE, checkFragLength=FALSE, countChimericFragments=FALSE, countMultiMappingReads=TRUE, fraction=TRUE.`

### 6. Construction of pseudo-bulk profiles

Throughout the manuscript, pseudo-bulk profiles refer to transcriptional profiles that are derived from the single cell reads by ignoring barcodes. This pipeline is depicted in **Figure 1 – figure supplement 2A**. By ignoring the R1 read of the single cell profile that contains the cellular barcode, we are effectively performing “bulk” RNA-sequencing using only the R2 read that aligns to a transcript in the sample. This collapses all cells to a single profile. We compared two different techniques to compute pseudo-bulk profiles. The *unfiltered* pseudo-bulk data is derived from counting raw reads aligned to the reference genome using the `STAR` tool (Dobin et al., 2013). The *filtered* pseudo-bulk dataset is obtained by first applying our sc-pipeline (`alevin-fry` followed by `EmptyDrops`) and then summing across all cells. The first is closer in spirit to true bulk (single read) profiling, while the second approach filters reads, cells and genes in the same manner as sc-analyses and therefore represents a middle point between bulk and sc- profiling. A comparison of *filtered* pseudo-bulk FCZ profiles at day 2 and 3 datasets indicated that the assay is robustly quantifying the expression of genes across different batches (**Figure 1 – figure supplement 3A**).

A comparison of bulk profiles versus *unfiltered* pseudo-bulk for the FCZ profiles at day 2 and 3 indicated that both methods identified all but 297 of the same genes. The missed genes tended to be expressed at low levels in the bulk profiles (**Figure 1 – figure supplement 3B**). Moreover, day 2 and day 3 pseudo-bulk profiles were significantly correlated with “true” bulk RNA-sequencing profiles (R ranges from 0.67 to 0.74; **Figure 1 – figure supplement 3C**). Here “true” bulk RNA-sequencing profiles were generated as described in **Methods 5**.

### 7. Whole genome DNA-sequencing

*C. albicans* populations were grown as described in **Methods 1**, although we did not apply a cell filtration step to remove filamentous cells. Preparation of genomic DNA used the MasterPure™ Yeast DNA Purification Kit (Lucigen # MPY80200) with a NextSeq500– 1flowcell mid output (130M fragments), 150 cycles pair-end reads (maximum 2×80 nt), yielding on average on average 23.9 million reads per sample (1 UT, FCZ at days 2, 3, 6, and 12). This gives an expected sequencing depth of 224, since the size of the *C. albicans* genome is ∼16Mb,

Raw reads were pre-processed with the sequence-grooming tool cutadapt (Martin, 2011) version 0.4.1 with quality trimming and filtering parameters: `--phred33 --length 36 -q 5 -- stringency 1 -e 0.1`. Analyses followed a previous study by Ford et al. (Ford et al., 2015) which also examined *C. albicans* complete genomes, although we used a more recent version A22 of the *C. albicans* SC5314 haplotype A genome for read mapping (downloaded from the Candida Genome Database; CGD; www.candidagenome.org). Briefly, reads were mapped using the BWA alignment tool (Li and Durbin, 2009) and RealignerTargetCreator and IndelRealigner from GATK (McKenna et al., 2010) were used for re-alignment. SNPs were detecting using Unified Genotyper (GATK version 1.4.14) using the same filtration criteria as reported in Ford et al. Determination of copy-number and loss-of-heterozygosity was perfmed using GATK with the strategy reported in Ford et al.

Our observed sequencing depth was just under 200. We identified approximately the same number of single nucleotide polymorphisms (N=3,304) and the same number of insertions/deletions (181/255 resp.) as a previous whole genome sequencing effort (Cuomo et al., 2019) using their bioinformatic pipeline. Although mutations were observed in some reads at some genomic loci, the samples were not enrichment for mutations that occurred more often than the rate of sequencing error which is 10^-3^ after correcting for multiple testing. This error rate is in line with estimates of the spontaneous mutation rate for *C. albicans* (Ene et al., 2018) (**Figure 2 – figure supplement 5**), together suggesting that the population is near isogenic without any significantly large subclones. Given that the error rate for copy number variants (e.g. loss, amplification) is lower than polymorphisms (Ene et al., 2018) and the duration of cell expansion before drug exposure (<2 days), it would be unlikely that spontaneous mutations explain the degree of heterogeneity that was observed 2-3 days post drug exposure.

### 8. Cell subpopulations, trajectory and signature analyses

To identify subpopulations of cells with similar gene expression patterns in an unbiased, unsupervised manner, we applied Leiden clustering (Traag et al., 2019) on the latent space generated by `scVI` (resolution of 0.4 for UT and FCZ day 3-6 analyses, and resolution of 0.5 for the analysis combining UT, FCZ day 2 and 3, CSP, and RAPA-treated cells). The lineage trajectory of day 3 and 6 FCZ-treated cells was performed using the R/Bioconductor `slingshot` package version 1.8.0 with default parameters (Street et al., 2018).

To investigate the key sources of variability across cell subpopulations, we curated 43 gene signatures related to microbial phenotypic diversity and drug tolerance including the cell cycle, TCA cycle, specific and general stress responses, metabolic pathways, amino acid biosynthesis, efflux pumps, and specific drug responses amongst others (Azadmanesh et al., 2017; Berman, 2006; Cote et al., 2009; Enjalbert et al., 2006, 2003; Gasch et al., 2000; Hardwick et al., 1999; Hinnebusch, 2005; Jackson et al., 2020; Natarajan et al., 2001; O’Duibhir et al., 2014; Pais et al., 2019; Sanglard, 2016; Sanglard et al., 2003b; Tsai et al., 2019; Yang et al., 2017) (**Figure 2 – table supplement 1**). In some cases, gene signatures from the literature were derived in other organisms and required orthology mappings to *C. albicans*.

Briefly, signatures of <u>cell cycle phases</u> were identified as transcriptional expression patterns in synchronous *C. albicans* populations (Berman, 2006; Cote et al., 2009) or were expert-curated and well-established cell cycle genes found in distinct clusters of *S. cerevisiae* single-cell profiles (Jackson et al., 2020). Signatures <u>specific to certain stresses</u> were found by transcriptional profiling of *C. albicans* challenged by temperature, osmotic and oxidative stress under conditions that permitted > 60% cell survival (Enjalbert et al., 2003) or in *C. glabrata* (Pais et al., 2019). We also curated more <u>general non-specific stress signatures</u> identified in *C.albicans* (Enjalbert et al., 2006) or *S. cerevisiae* (Gasch, 2007; Gasch et al., 2017; Tsai et al., 2019). Previous studies established the existence of a ubiquitous Environmental Stress Response (ESR) in *S. cerevisiae* which is deregulated in response to many different environmental perturbations (Gasch, 2007; Gasch et al., 2000). The ESR is divided into the induced (iESR) and repressed (rESR) subcomponents. The iESR is characterized by overexpression of heat-shock and oxidative stress genes in addition to genes involved in central carbohydrate metabolism and energy generation (Gasch, 2007; Gasch et al., 2000). Previous studies have noted a complex, intricate relationship between the ESR and cell cycle phase (O’Duibhir et al., 2014; Regenberg et al., 2006). Some investigations suggested a smaller core ESR in *C. albicans* (Enjalbert et al., 2006) which may have evolved due to the unique host environment with the need to grow with different substrates (Brown et al., 2014). Curated <u>metabolic signatures</u> include lowly expressed glycolytic genes and highly expressed TCA cycle genes identified during diauxic shift or rapamycin treatment in yeast (DeRisi et al., 1997; Hardwick et al., 1999) as well as GCN4-driven amino-acid biosynthesis, another well defined metabolic signature conserved between *S. cerevisiae* and *C.albicans* (Hinnebusch, 2005; Natarajan et al., 2001).

The signature analyses start with the `VISION` tool which estimates a signature score for every cell (DeTomaso et al., 2019) using the batch-adjusted normalized counts returned by `scVI`’s model. The distribution of individual scores for cells classified in each cluster can be depicted using the empirical cumulative distribution function. These distributions were further compared using the Kolmogorov-Smirnov test. We then used the median to summarize signatures scores of cells within each cluster and selected signatures which were the most variable across clusters (sd > 0.01 for UT analyses and sd > 0.05 for the analysis combining UT, FCZ- day 2 and 3, CSP-, and RAPA-treated cells or combining FCZ-treated cells at day 3 and 6 ; **Figure 2 – table supplement 1).** Heatmaps were used to depict the median scores (z-score, color bar) of the selected signatures for each cluster.

We further investigated the expression of individual genes from signatures expressed in UT cells. To do so, we first used MAGIC (van Dijk et al., 2018) for imputation of missing expression measurements. We computed gene correlation matrices for each signature to verify good concordance of expression across cells. We used the expression of the most variable genes (sd > 0.03) to order UT cells from high to low mean gene rank. We plotted heatmaps to present the expression (z-score, colorbar) of genes positively correlated with the cell ordering. The mean gene rank orderings were used for downstream analyses of UT cells allowing 3-way comparisons of signatures expressed in individual cells (**Figure 2E,F**).

### 9. Differential gene expression (DGE) analysis

#### Single-cell DGE

`scVI`’s model allows us to approximate the posterior probability of the alternative hypotheses (genes are different) and that of the null hypotheses through repeated sampling from the variational distribution, thus obtaining a low variance estimate of their ratio (i.e., Bayes factor). We used this approach to identify genes differentially expressed in the combined and individual comet clusters compared to the other clusters (**Figure 3 – figure supplement 2A,B**).

#### Pseudo-bulk DGE and GO enrichment analysis

We also conducted pseudo-bulk differential gene expression analyses which allow for a dramatic reduction in the number of zeros in the data by aggregating cells within each replicate. This approach has been found to achieve the highest fidelity to the experimental ground truths significantly reducing the risk of false discoveries (Squair et al., 2021). As described in **Methods 6**, *filtered* pseudo-bulk profiles were obtained by summing counts of selected cells within each replicate. In order to use `DESeq2` (Love et al., 2014), a standard R/Biocoductor package used for differential analysis of count data, we require at least two replicates within each group of comparison. If only one replicate was available (e.g., CSP, RAPA, FCZ at day 6), we partitioned the selected cells of a single replicate into two groups randomly before forming pseudo-bulk profiles. We then used `DESeq2` default parameters to perform DE analysis and selected genes with Benjamini-Hochberg FDR < 0.1 (Benjamini and Hochberg, 1995). Models were adjusted for batch affects where relevant.

Finally, we identified enrichment of biological processes in lists of significantly over- and under-expressed genes using the R/Bioconductor `ViSEAGO` package (Brionne et al., 2019). `ViSEAGO` includes all algorithms developed in the R/Bioconductor `topGO` package including the weight01 fisher test that takes into account the topology of the Gene Ontology (GO) graph (Alexa et al., 2006). Biological processes with weight01 p-value < 0.01 were defined as significantly enriched in the gene list.

To further compare GO enrichments of gene lists associated with the α and β populations at day 3 and 6, we first used Wang’s graph-based method implemented in `ViSEAGO` to compute semantic similarity between GO terms. Groups of enriched GO terms were then produced by cutting the dendrogram in a static or dynamic mode developed in the `dynamicTreeCut` R package (Langfelder et al., 2008).

### 10. Cell imaging

To validate the subpopulation structure identified by the sc-transcriptomics, we choose markers representative of distinct clusters. Cells were transformed with *GFP* and *RFP* fusion constructs for *HSP70* and *TTR1* marker genes respectively using a CRISPR/Cas9 protocol (Min et al., 2016) with primers described in **Figure 2 – table supplement 2**. Strain SN76(*his1Δ/his1Δ, arg4Δ/arg4Δ, ura3Δ/ura3Δ*) was chosen for gene tagging, since it is a derivative strain of SC5314 but with multiple auxotrophic markers. These markers allow for convenient selection of successfully transformed cells.

Benchling (https://benchling.com) was used to design the sgRNAs. We followed the CRISPR/Cas9 protocol with the plasmid pV1093 from Min et al. (Min et al., 2016). This includes two PCR reactions to fuse the SNR52 promoter to the sgRNA scaffold and terminator. The third PCR reaction amplifies the final sgRNA cassettes. Two different plasmids pENO1-iRFP-NATr (Plasmid #129731, Addgene Inc) and pFA-GFP-HIS1 were used to design the repair segments. The construction of the Cas9 cassette proceeded as per Min et al. Amplification of the Cas9 cassette with PCR used the following schedule: 98°C for 3 minutes, 98°C for 30 seconds, 63°C for 30 seconds, 72°C for 5 minutes and 30 seconds. Steps 2 to 4 have been repeated for 34 rounds followed by 72°C for 10 minutes and finally the reaction finished in 4°C. The repair DNA must be amplified with the designed primers (**Figure 2 – table supplement 2**) in 8-12 PCR tubes with 0.1μl plasmid (500ng/ml), 2.5μl forward primer, 2.5μl reverse primer, 1μl 10mM dNTP, 33.65μl nuclease free water, 10μl 5X HF PCR buffer and 0.25μl phusion polymerase in each tube.

Cells that were successfully transformed were grown and harvested in a manner identical to that used for the single cell experiments (**Methods 1**). At time of microscopy, cells were collected, washed with H2O and transferred to minimum media to minimize the background noise from normal YPD media. Afterwards, cells were mounted onto the uSlide and imaged with a Leica DM6000 microscope at 1000x (∼50 images/time point; ∼50 cells/image).

## Author Contributions

SM led the development of the lab work associated with the project and VD led the data science/computational biology components of the project. Both contributed to the analysis of the data and preparation of the manuscript with early contributions from VB. AC and RO provided assistance with the preparation of cultures and design of the experiments. SK provided assistance with the DROP-seq assay. SS contributed to the microscopy aspects of the project. LX, MW, JB, VD and MH provided assistance with writing the manuscript. MW provided laboratory support and MH funded the project.

## Competing Interests

The authors declare that they have no conflicts of interest.

## Acknowledgements

We thank L. Pachter and the organizers of BIRS workshop #17w5134 (Oaxaca, Mexico) for several conversations that motivated this effort especially with respect to affordable open biotechnology. We thank C.A. Jackson and D. Gresham for early access to their protocol, C. Law for assistance with the microscopy, A. Villani for several nice optimizations, and members of the McCarroll lab for their careful advice and outstanding responsiveness.

## Data Availability and Reproducibility

Python/R code and data required for reproducibility is available through the Open Science Foundation (OSF) repository https://osf.io/5tpk3/ and associated github repository https://github.com/vdumeaux/sc-candida_paper. The raw and processed single-cell transcriptome and bulk RNA-seq is also available through NCBI’s Gene Expression Omnibus with accession number GSE204903.

## Appendix 1. Drug diffusion assays with FCZ treated cells at days 1-6

Diffusion disk assays (DDAs) provide a means to measure tolerance and resistance of a fungal population to a drug (Gerstein et al., 2016). In our case, we use these assays to measure tolerance and resistance of *C. albicans* in response to FCZ. Towards this end, two SC5314 colonies were transferred from YPD agar plates to 5 ml YPD liquid media and incubated at 30°C overnight. Cells were washed the next day with PBS 1X twice and an aliquot of 10^6^ cells/ml was transferred into an eppendorf tube. 100 ul of this suspension was plated on YPD agar and spread evenly with a cell spreader. A fluconazole disk (BioRad #62802) was placed at the center of the plate and incubated at 30°C for 48 hours. This was repeated three times (replicates 1-3) for a total of two biological replicates (experiments 1-2).

Images were captured every 24 hours for 6 days (**Appendix 1 – figure 1**). **Appendix 1 figure 2** describes how the radius (RAD) and the Fraction of Growth (FoG) are estimated from these images to provide measures of drug resistance and tolerance respectively. We use the `diskimageR` R package to estimate the intensity of 72 lines (in 5° increments around the circle) from the disk edge to the rim of the plate for each replicate across 6 days post-exposure (**Appendix 1 figures 3).** A Kruskal-Wallis χ^2^ test was used to test whether there was a difference in the mean of either the FoG or RAD between day 1 and the remaining time-points versus a null hypothesis of no difference.

**Appendix 1 Figure 1.**
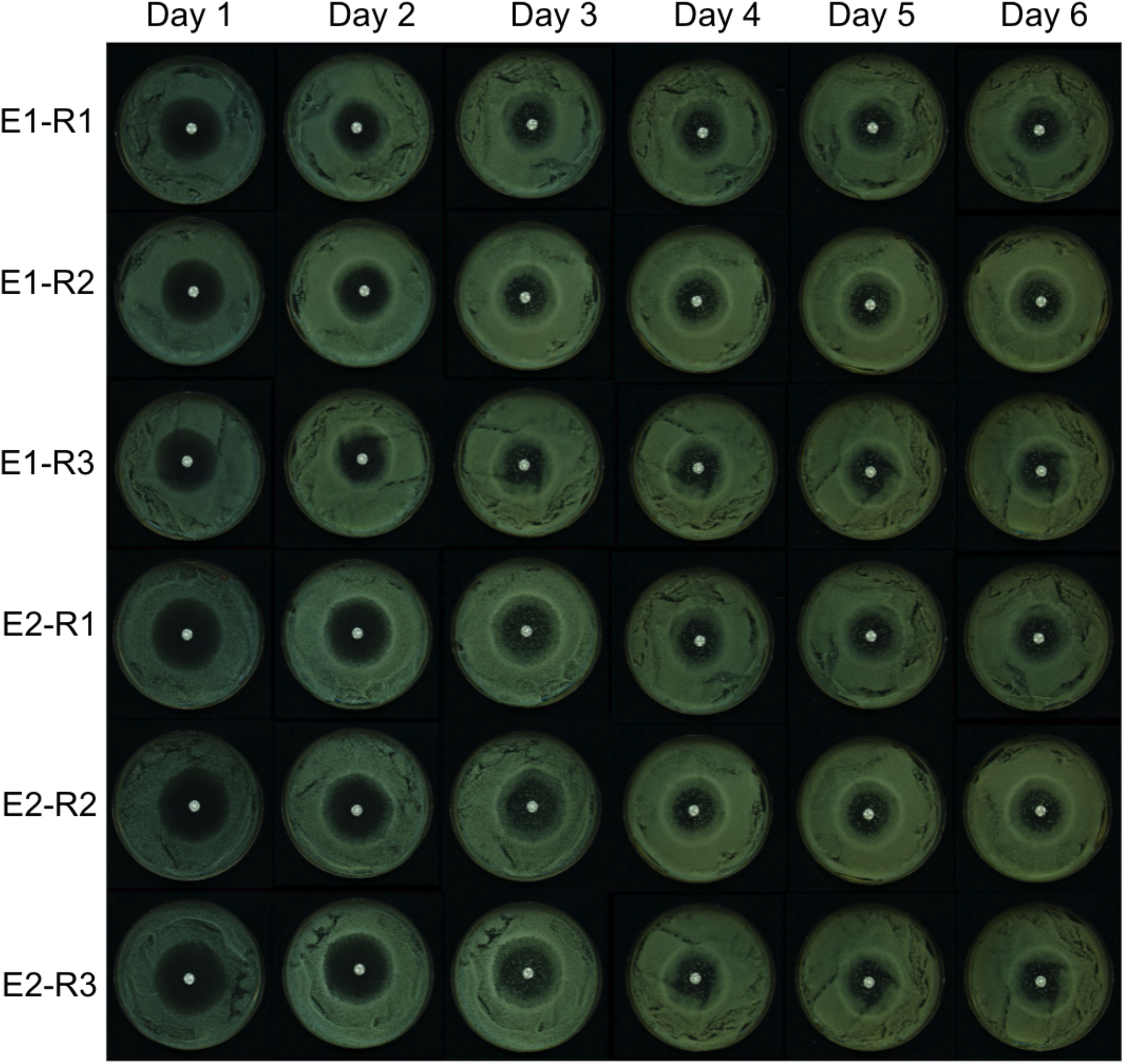
Fluconazole disk assay results across six days. Two colonies (biological replicates E1 and E2) of *C. albicans* SC5314 were grown in YPD and spread on three plates (technical replicates R1, R2 and R3). The bright white spot at the center of the plate is the FCZ diffusion disk.

**Appendix 1 Figure 2.**
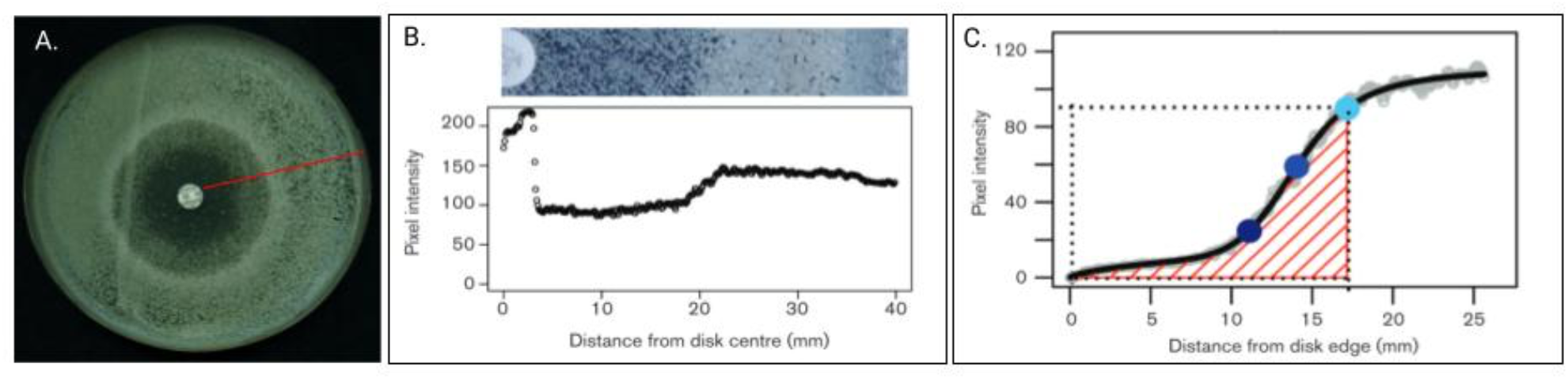
**(A)** A *C. albicans* colony with FCZ disk in the middle of plate. The red radial line represents one of 72 measurements taken every 5°. **(B)** Top is a blown up and restricted image of the red line from (A) and bottom represents the pixel intensities from 0 (edge of disk) to 40 (edge of plate). **(C)** Grey curves are from the 72 measured radial lines. Black represents the average of these 72 measurements per mm from disk edge (x-axis). Light blue dot is the RAD_20_, defined as the point where there is a 20% reduction in growth. Middle blue dot is the RAD_50_ and dark blue represents the RAD_80_. The FoG_20_ is defined as the fraction of the area under the black line from 0 to the RAD_20_ (that is, the amount of growth observed) divided by the total potential growth (delimited by dotted line). The FoG_50_ and FoG_80_ are defined analogously (adapted from Gerstein et al. 2016.)

**Appendix 1 Figure 3.**
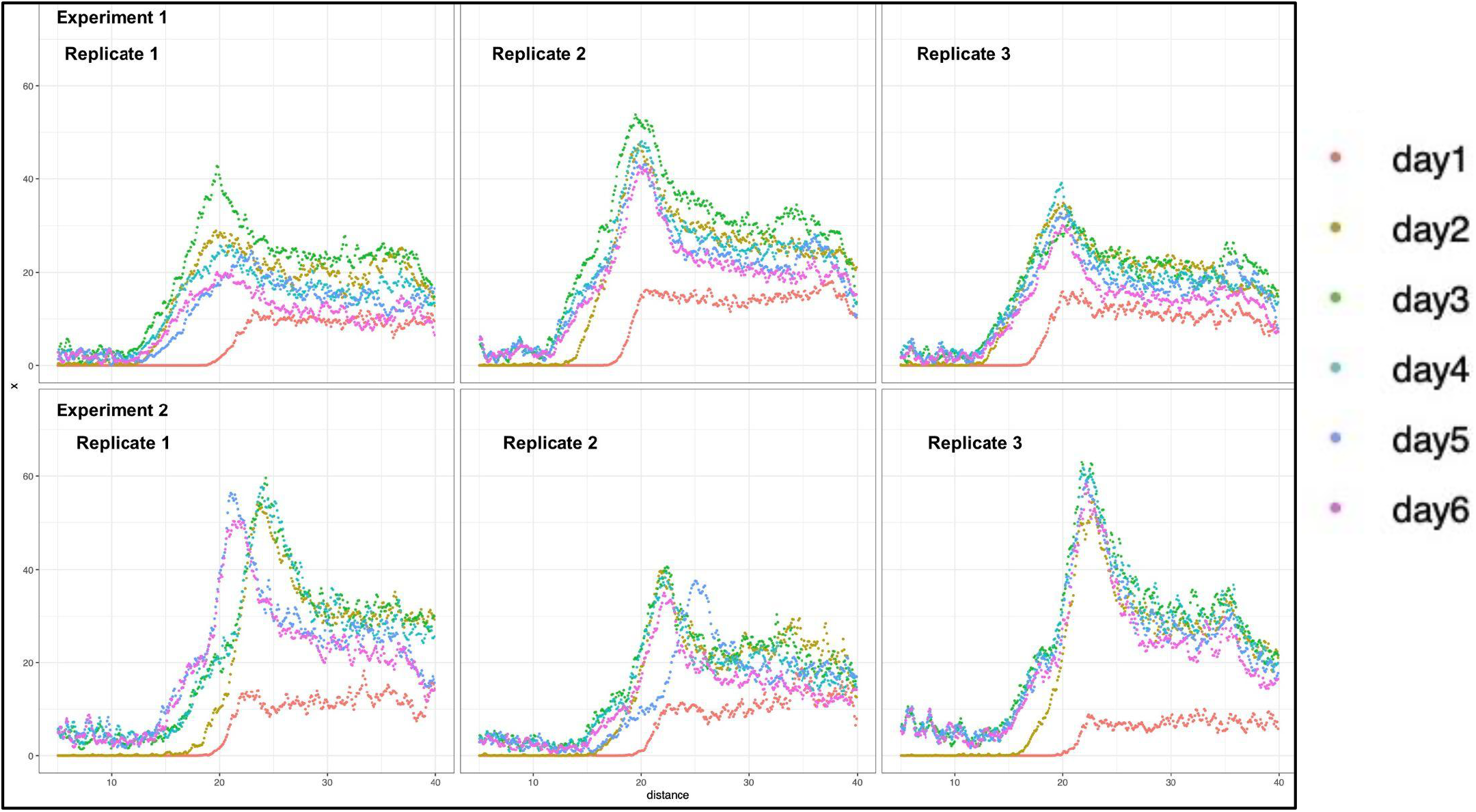
Two biological (independent *C. albicans* SC534 colonies) and three technical replicates of FCZ DDA experiments. Here the x-axis is distance from the disk in millimeters (0-40mm) and the y-axis is intensity where higher intensity indicates more *C. albicans* growth. The red curve represents 1 day post-exposure; we can see that the “flat” Zone of Inhibition ends at approximately 20mm on average (RAD_80_). Then the positive “steep” slope continues until 23mm on average (RAD_20_). This 3mm region is the tolerance zone. In contrast, already at day 2 the “flat” Zone of Inhibition ends earlier at 15mm on average, and the slope increases steadily until ∼21mm on average, defining a tolerance zone of 6mm, a highly significant increase in tolerance (p < 0.001, Kruskal-Wallis χ^2^). There is a slight but statistically reduction in the RAD_20_, suggesting that there has been a small increase in resistance (p < 0.05, Kruskal-Wallis χ^2^). However, we can see that the vast majority of the colony is not resistant. The same trend is observed for Days 3-6 but note that the intensity is slightly lower, likely reflecting the fact that FCZ has dissipated by these later time points.

---

**Figure 1 - figure supplement 1.**
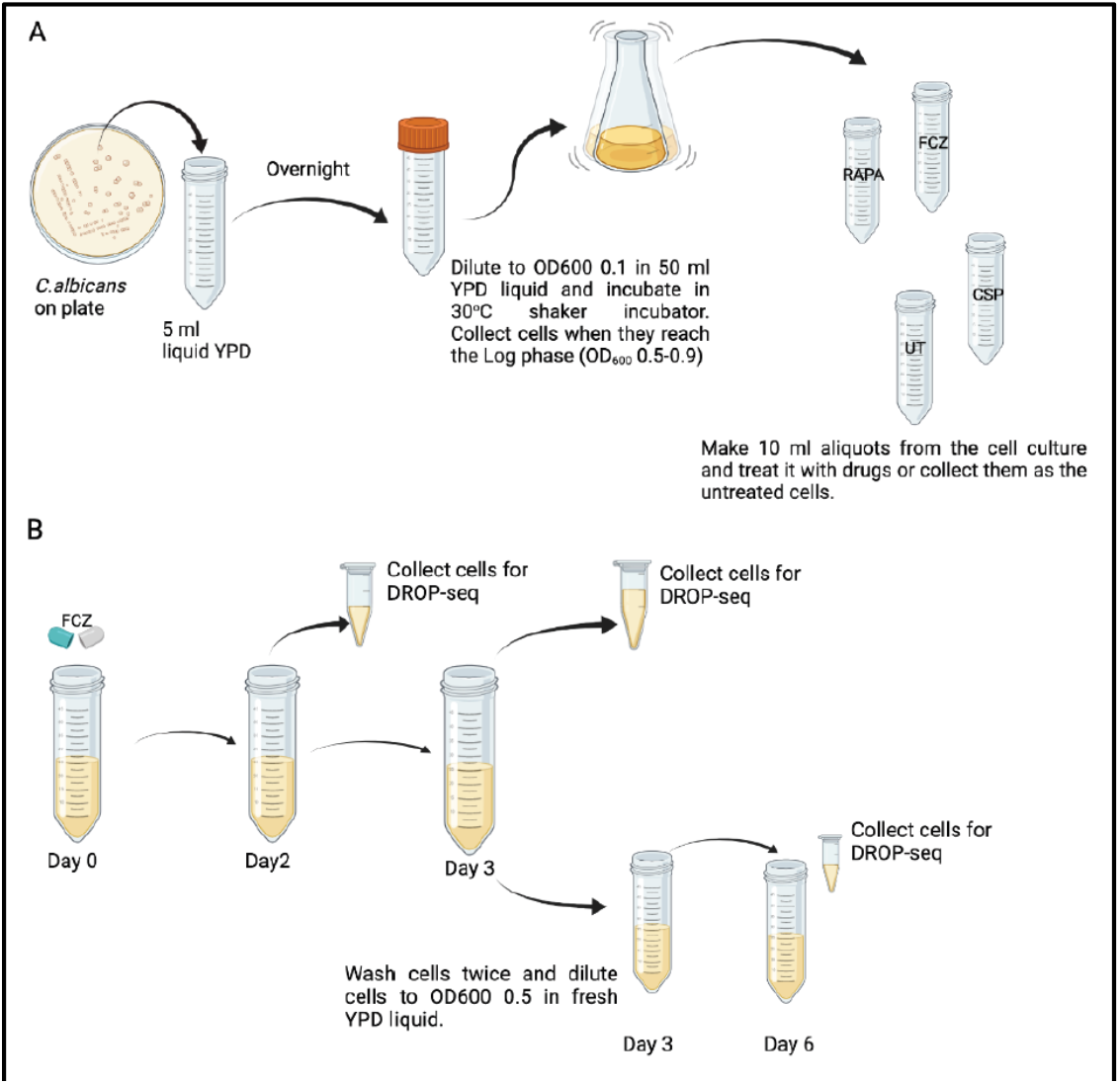
**(A)** Schematic of cell growth: Cells untreated (UT) or treated with fluconazole (FCZ), caspofungin (CSP) and rapamycin (RAPA). **(B)** Schematic of sample collection for samples that underwent FCZ exposure (collected on days 2 and 3) and recovery in YPD (collected on day 6).

**Figure 1 - figure supplement 2.**
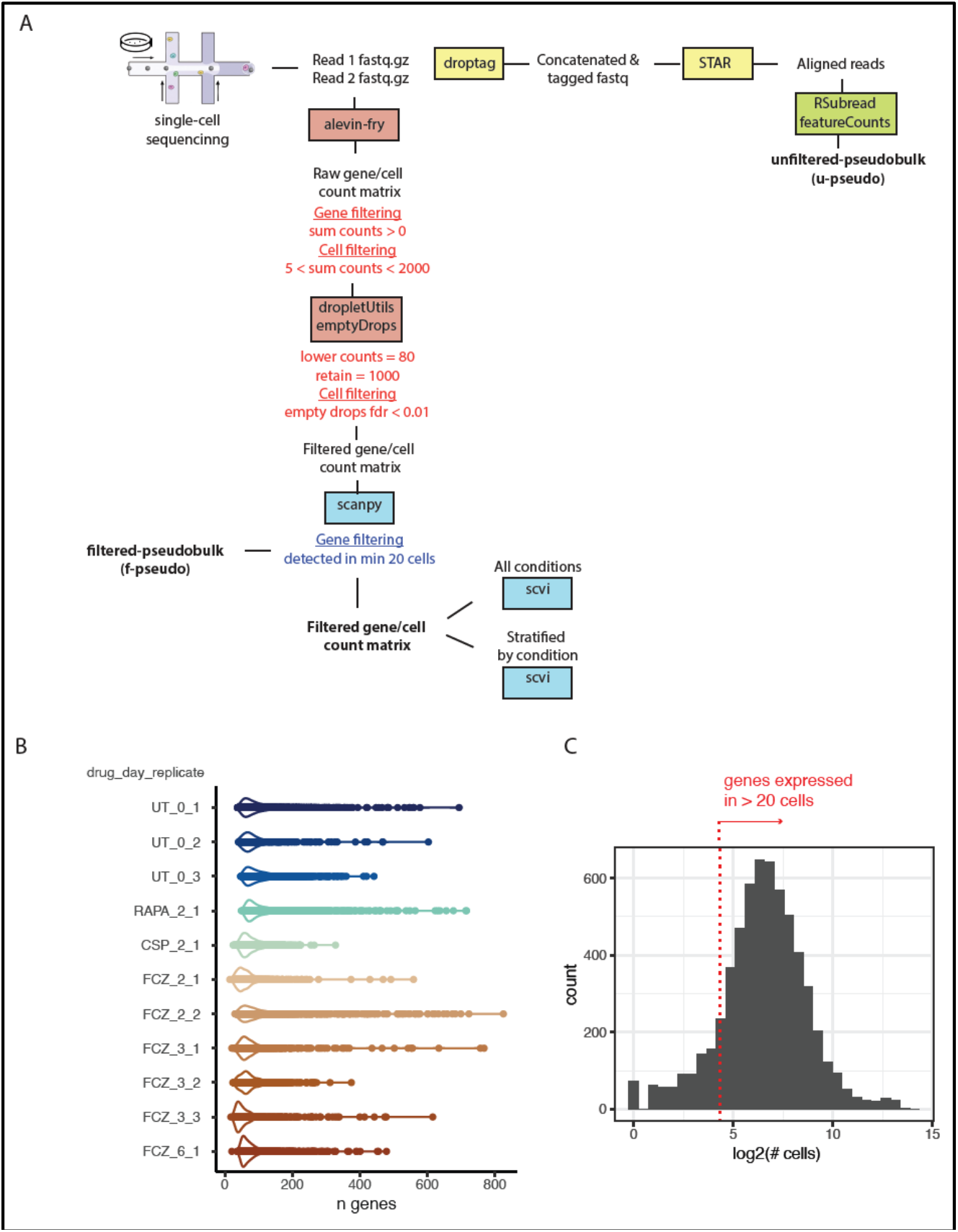
**(A)** The bioinformatics pipeline that begins with the Read1 (left) and R2 (right) fastq files (**Methods 4**). Sc- sequence data was processed using the Alevin-Fry (He et al., 2021) pipeline based on a reference index covering the spliced transcriptome . Cells were labeled as good quality if their profiles significantly deviated from the ambient RNA pool (FDR < 0.01) following the EmptyDrops approach. We then filter genes that were detected in less than 20 cells. Pseudo-bulk profiles were obtained by either summing counts across selected cells and genes (filtered pseudo-bulk) or by mapping reads to the reference genome but ignoring cell barcodes (unfiltered pseudo- bulk) (**Methods 6**). **(B)** Violin plots with data points describing the number of genes detected in each high quality cell for each sample (drug_day_replicate) **(C)** Histogram depicting the number of cells in which a gene is expressed. We selected genes that were expressed in at least 20 cells across all samples (red dotted line).

**Figure 1 - figure supplement 3.**
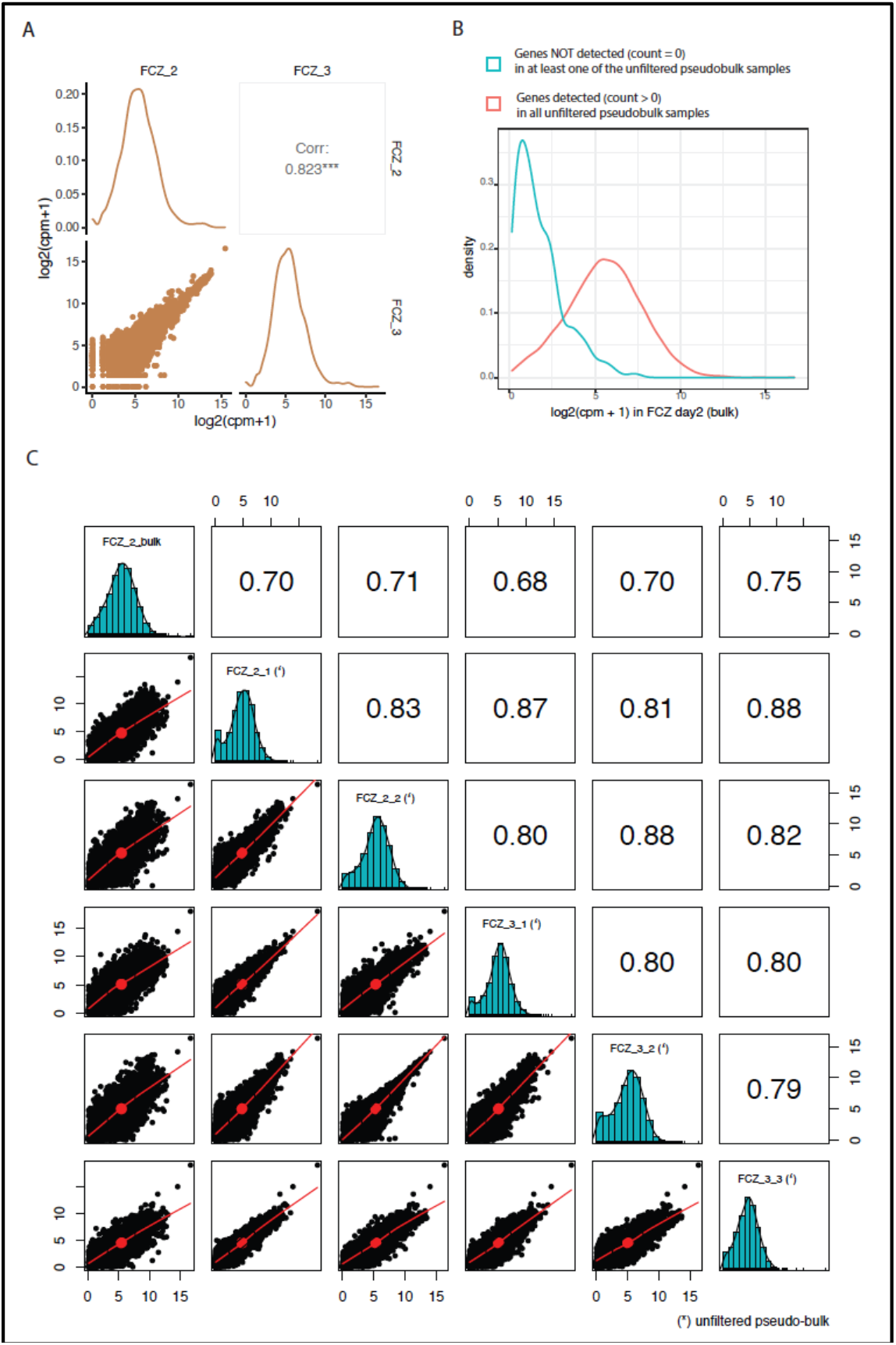
**(A)** The diagonal of the matrix shows the distribution of gene counts in single-cell profiles of FCZ-treated cells at day 2 (FCZ_2) and day 3 (FCZ_3). Counts for each high quality cell in both day 2 and day 3 replicates were summed and normalized (filtered pseudo-bulk, see figure 1 – figure supplement 2A; Counts Per Million, CPM). The upper right and lower left quadrants report the Pearson correlation and scatter plot depicting how genes are expressed in the two filtered pseudo-bulk samples. **(B)** The density plots compare the normalized counts in the bulk profile of FCZ- treated cells at day 2 of genes that were consistently detected in every FCZ (unfiltered) pseudo-bulk (red) versus genes that were not consistently identified (blue). Unfiltered pseudo- bulk samples were obtained by aligning single- end reads ignoring the paired-read containing the cell barcode for each replicate (figure 1 – figure supplement 2A). **(C)** The diagonal of the matrix shows the distribution of normalized gene counts, measured as the log_2_(CPM+1) in each FCZ-treated sample (bulk or (*)unfiltered pseudo- bulk). The values above the diagonal report the Pearson correlations between pairs of sample, and the scatter plots below the diagonal depict normalized log_2_(CPM+1) expression of genes in each pair of samples (bulk or (*) unfiltered pseudo-bulk).

**Figure 2 - figure supplement 1.**
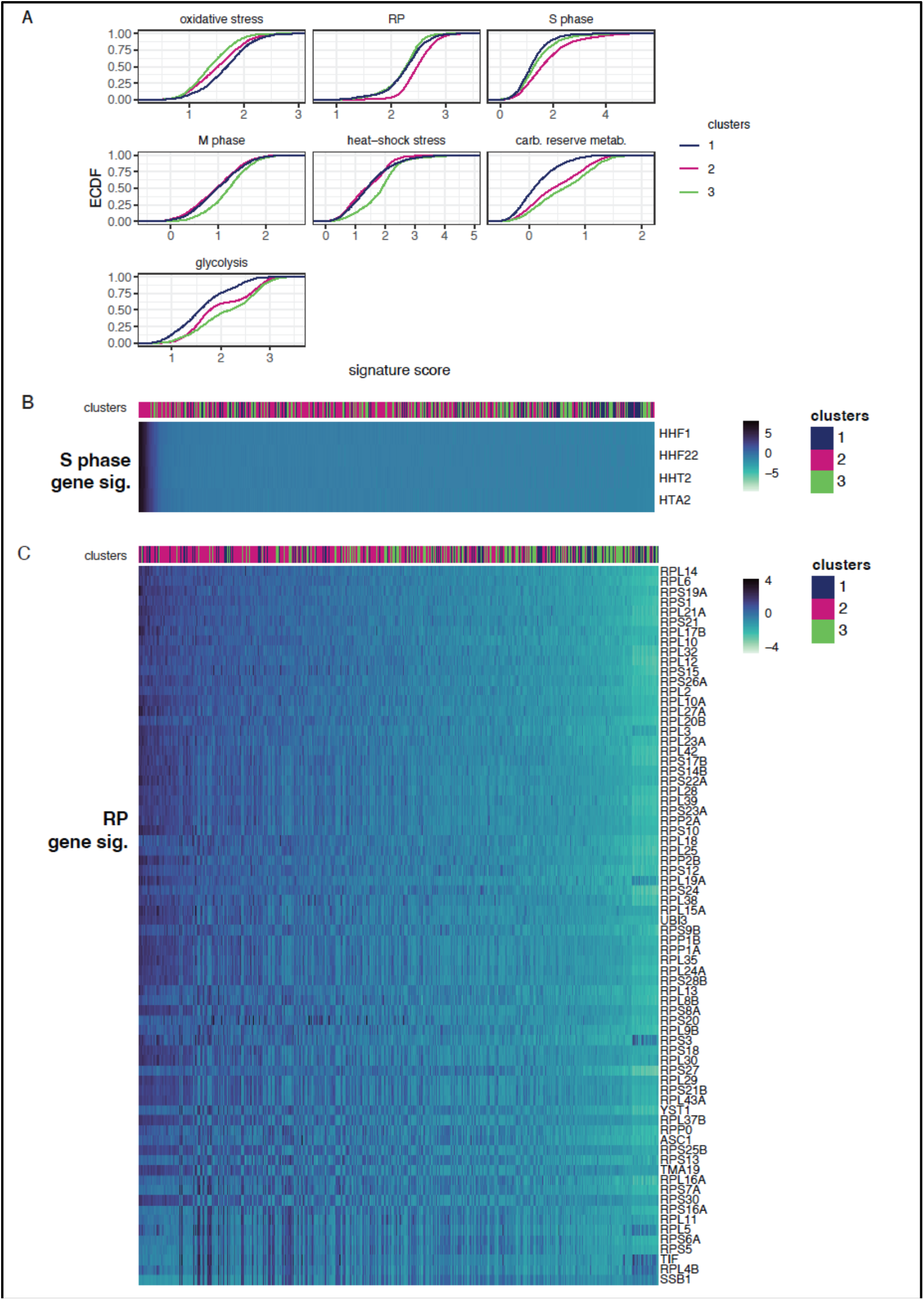
**(A)** Empirical cumulative distribution (ECDF) of signature scores for each cell in the distinct clusters (color) presented in Figure 2A. The y-axis provides the proportion of cells in each cluster whose scores fall below a certain value (x- axis). **(B)** For the S/G2 signature, and **(C)** for the ribosomal protein (RP) signature, the heatmap depicts the expression of selected genes (rows) across cells (columns). We used MAGIC to impute expression levels (z- score, color bar). Cells are linearly ordered by the overall magnitude of gene expression (mean gene rank). “Clusters” (top) indicate in which cluster the cell is assigned.

**Figure 2 - figure supplement 2.**
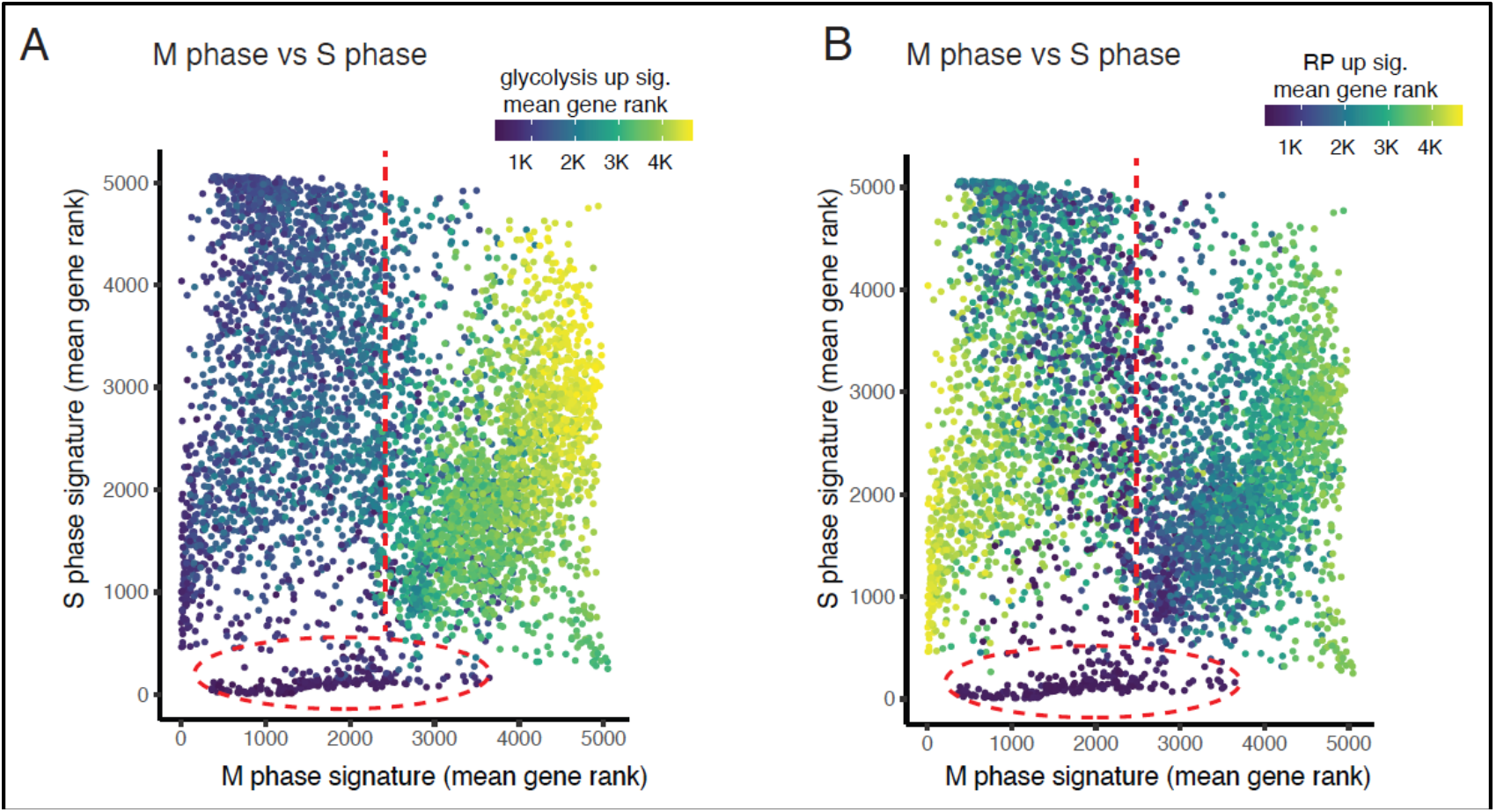
For each cell (point), the overall magnitude of gene expression (mean gene rank) of the M and S phase signatures were plotted. Color bars indicate the magnitude of gene expression (mean gene rank) of the **(A)** glycolysis, **(B)** RP-coding genes signatures.

**Figure 2 - figure supplement 3.**
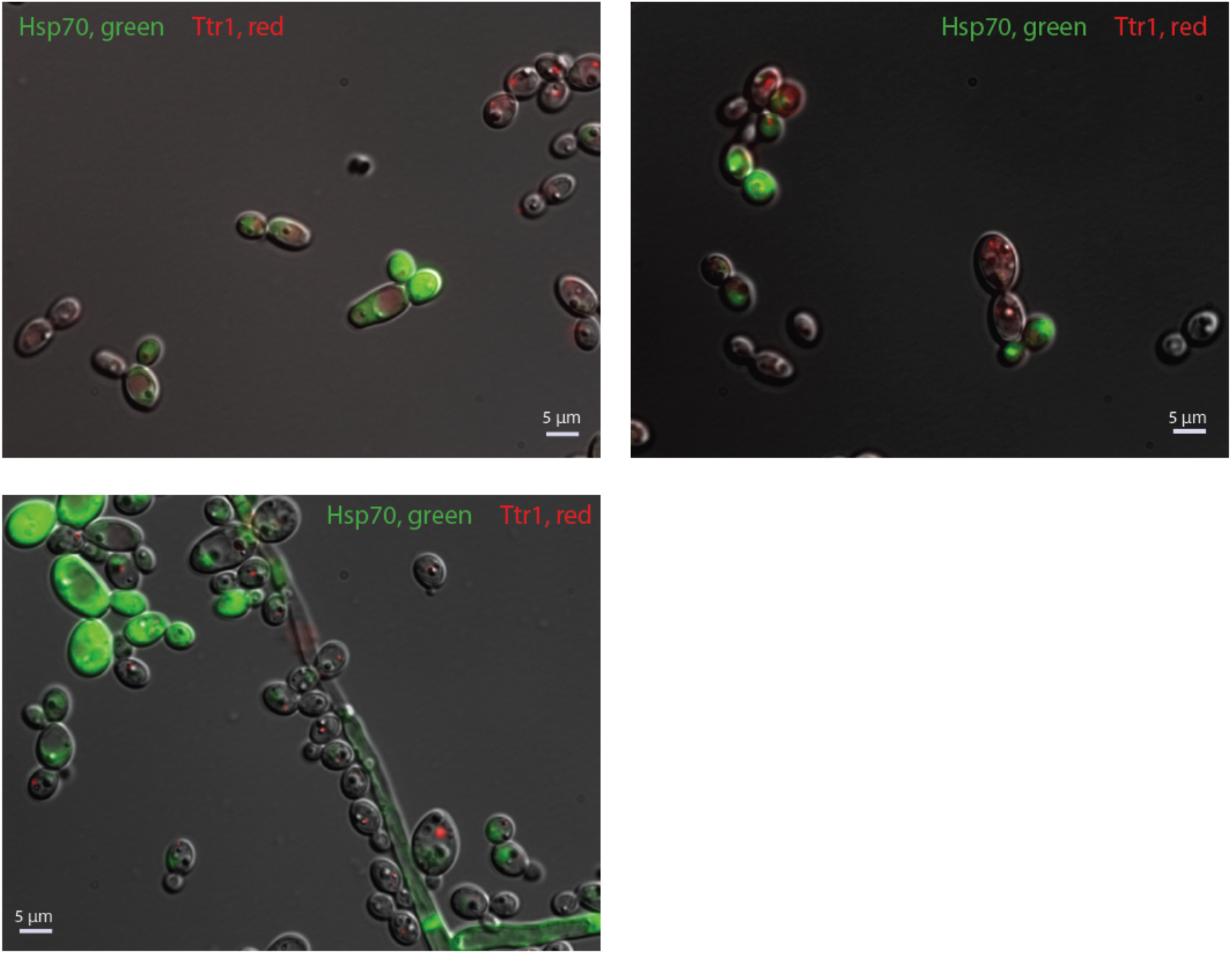
Examples of microscopy images of *C. albicans* cells harboring both a GFP-tagged HSP70 and an RFP-tagged TTR1. We observe mutual exclusivity in expression: cells tend to express either HSP70 or TTR1, but not both.

**Figure 2 - figure supplement 4.**
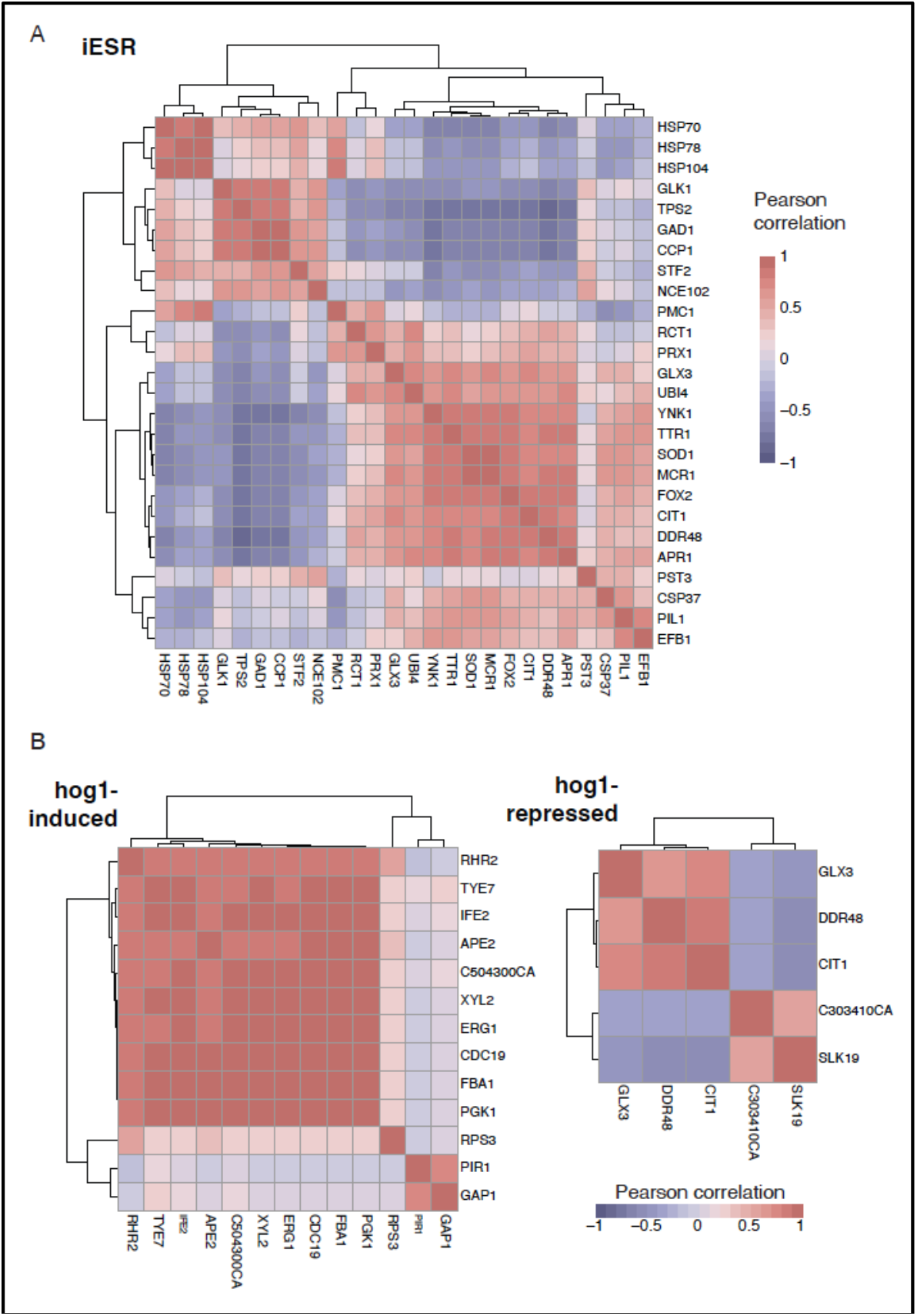
Correlation matrices depicting co-expression of genes **(A)** from the induced Environmental Stress Response (iESR) (Gasch, 2007; Gasch et al., 2000) and **(B)** induced (left) and repressed (right) in the Hog1-driven stress signature (Enjalbert et al., 2006) in UT cells.

**Figure 2 - figure supplement 5.**
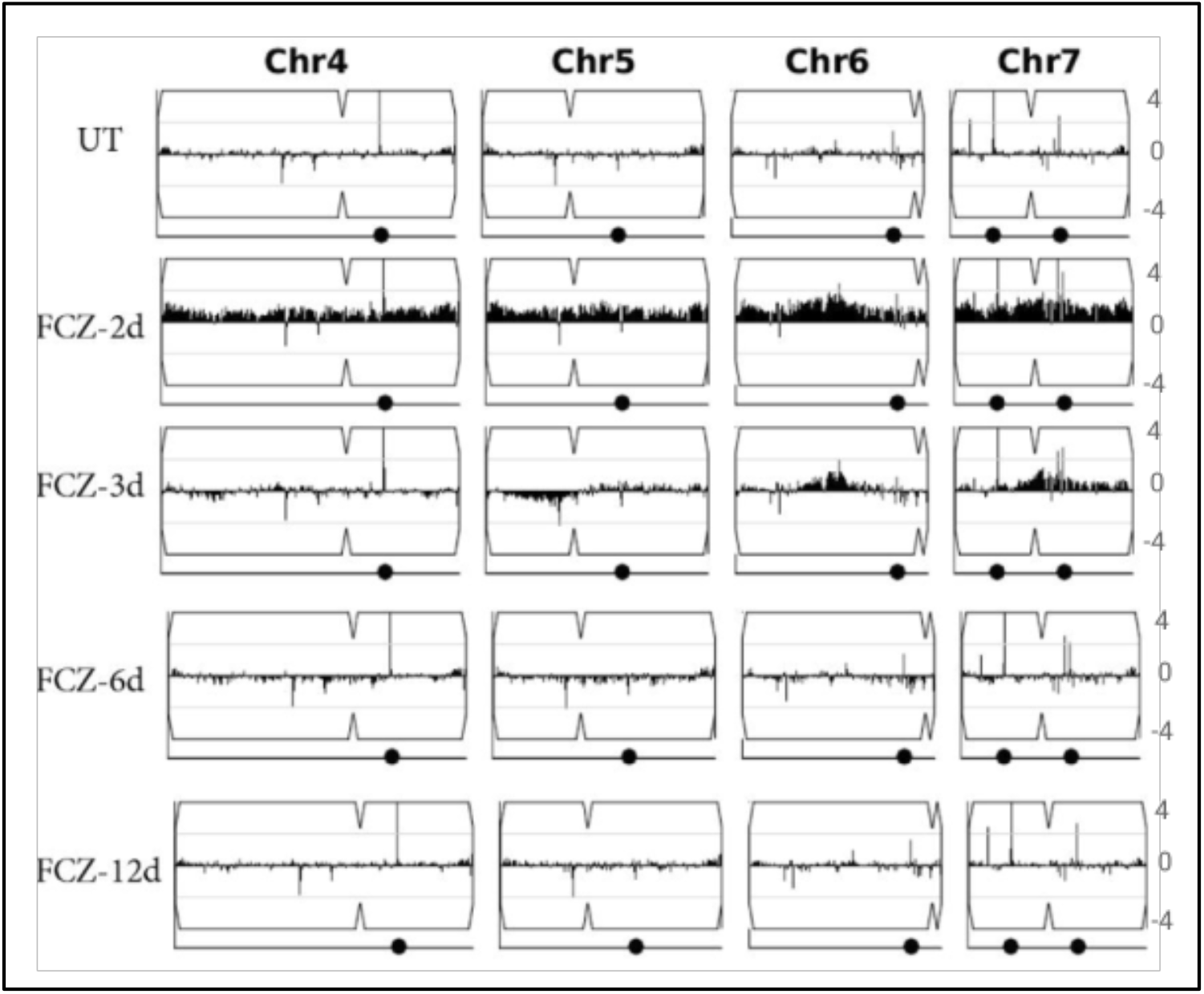
A summary of chromosomal instability across four C. albicans chromosomes at five time points (UT followed by 2, 3, 6 and 12 day time points). Here the axis traverses the four chromosomes. The amplitude of each point along the y-axis normalized to represent the deviation from a normal diploid genome with points above and below the line representing amplification and deletion respectively. Here amplitude is the log2 of the number of reads obtained from whole genome DNA sequencing.

**Figure 3 - figure supplement 1.**
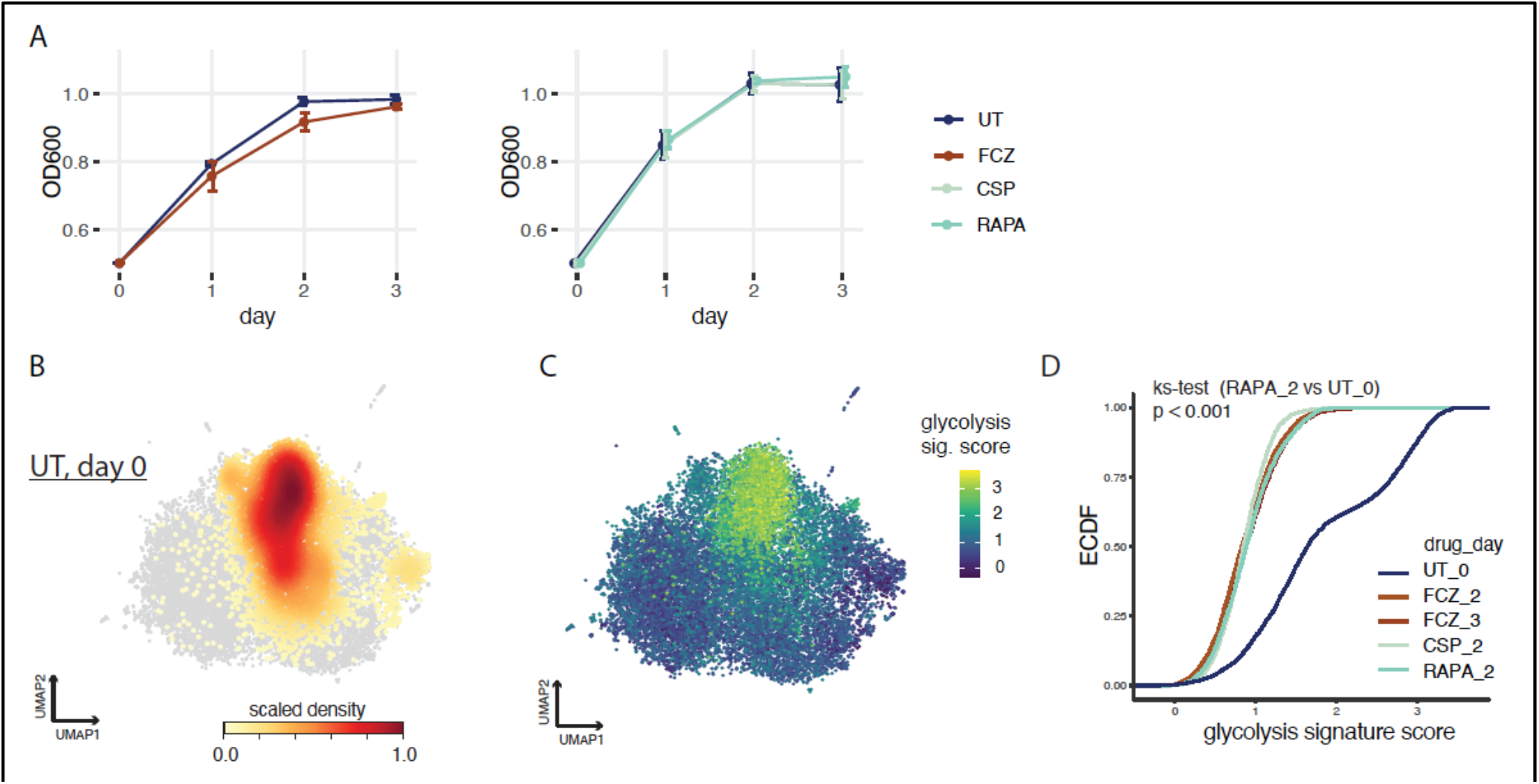
**(A)** Growth curves for untreated (UT) cells and cells treated with the three antifungals (FCZ, fluconazole; CSP, caspofungin; RAPA, rapamycin) as measured by OC_600_ across days. **(B,C)** UMAP embedding from figure 3B. Color bar depicts: **(B)** density of UT cells and **(C)** expression score for the glycolysis signature. **(D)** Empirical cumulative distribution (ECDF) of the glycolysis signature scores for each cell in the different conditions (drug_day, color). ks-test, Kolmogorov-Smirnoff test.

**Figure 3 - figure supplement 2.**
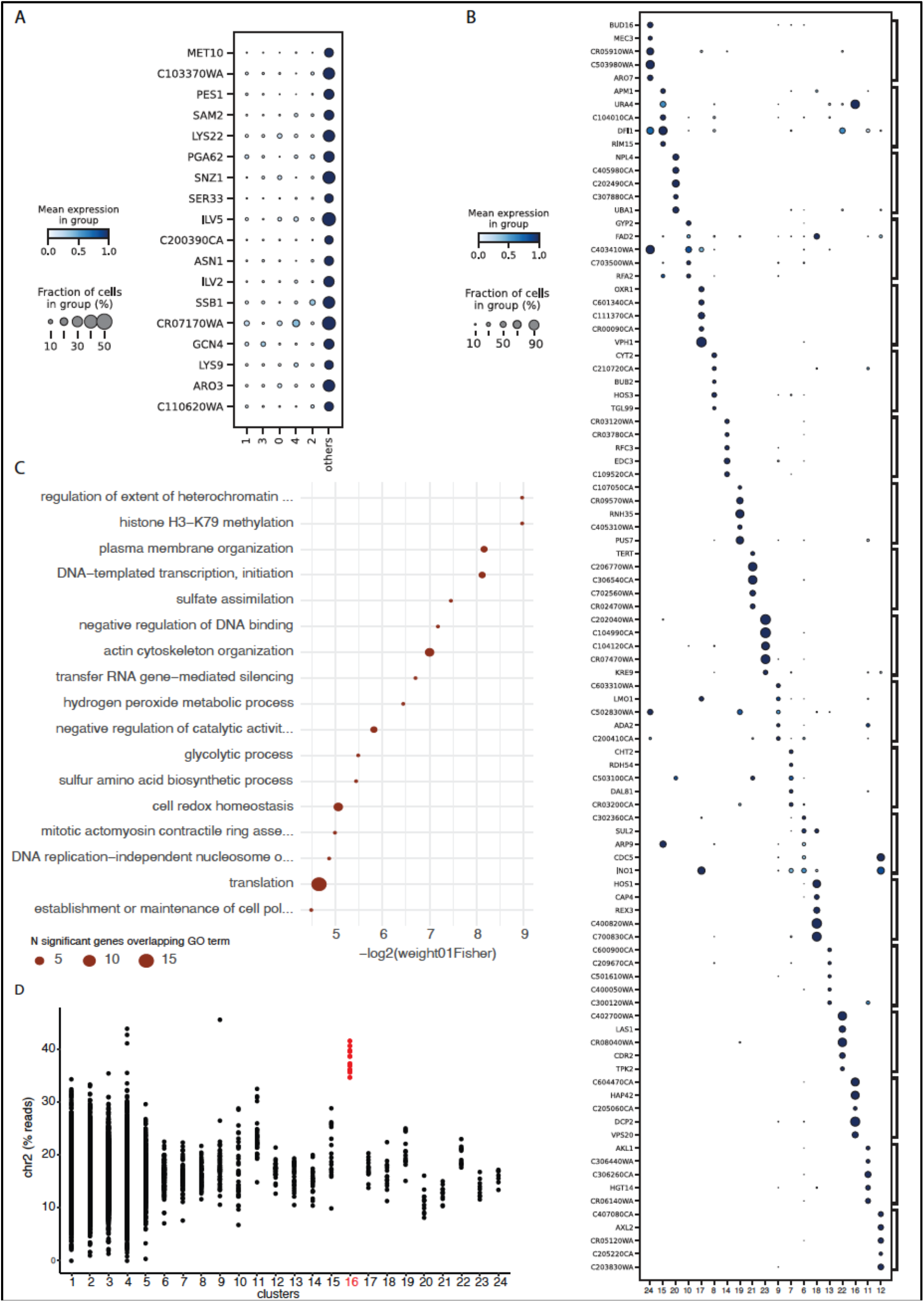
**(A)**Genes overexpressed in the collection of 19 comets compared to other cells in main clusters (Bayes factor > 2.5 and proportion of non-zero value > 0.2). Color- bar indicates the mean expression across the cells in each cluster. The size of the dot indicates the proportion of cells which express that gene within each cluster. (**B**) similar plot than (A) for genes differentially expressed in each comet cluster compared to others (Bayes factor > 3 and proportion of non-zero value > 0.2). **(C)** GO enrichment for genes identified as differentially expressed in comets compared to other clusters using pseudo-bulk differential expression analysis (60 genes; FDR < 0.1 ; Figure 3 – table supplement 1A,B).The size of the dot is proportional to the number of genes in the list which overlap with the corresponding GO term. **(D)** Percent of reads assigned to genes in chromosome 2 for each cell classified in the 24 distinct clusters.

**Figure 3 - figure supplement 3.**
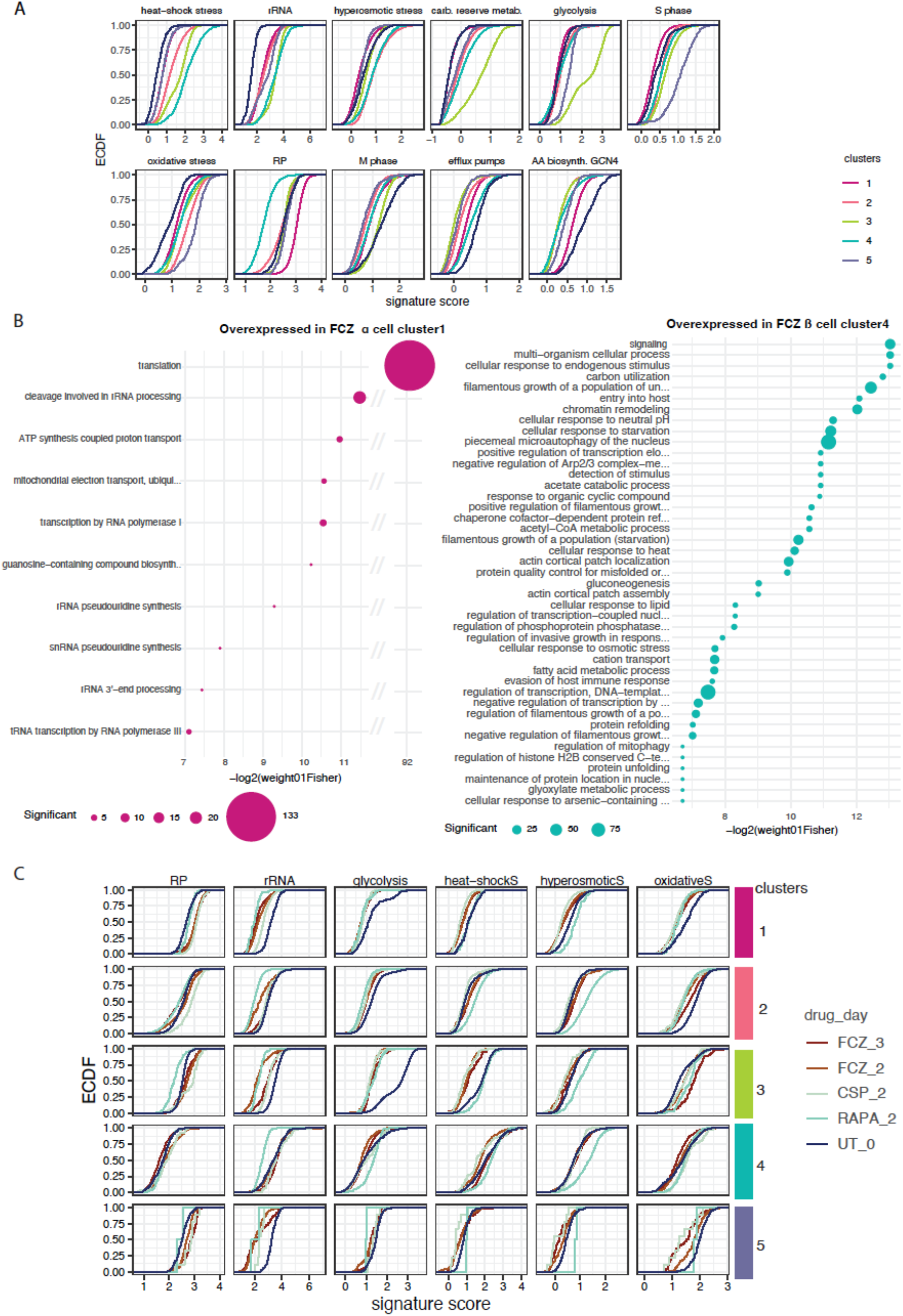
**(A)** Empirical cumulative distribution (ECDF) of signature scores for each cell in the distinct clusters (color) presented in Figure 3B. The y- axis provides the proportion of cells in each cluster whose scores fall below a certain value (x-axis). **(B)** GO enrichment for genes identified as differentially expressed in FCZ α cell cluster 1 compared to FCZ β cell cluster 4 using pseudo-bulk differential expression analysis (N = 223 and 463 genes overexpressed in α (left) and β (right) cells, respectively ; FDR <0.1). The size of the dot is proportional to the number of genes in the list which overlap with the corresponding GO term. **(C)** Similar to plot (A) but distributions are given for signature scores in each cell from different conditions (colored curves) and classified in distinct clusters (facet rows).

**Figure 3 - figure supplement 4.**
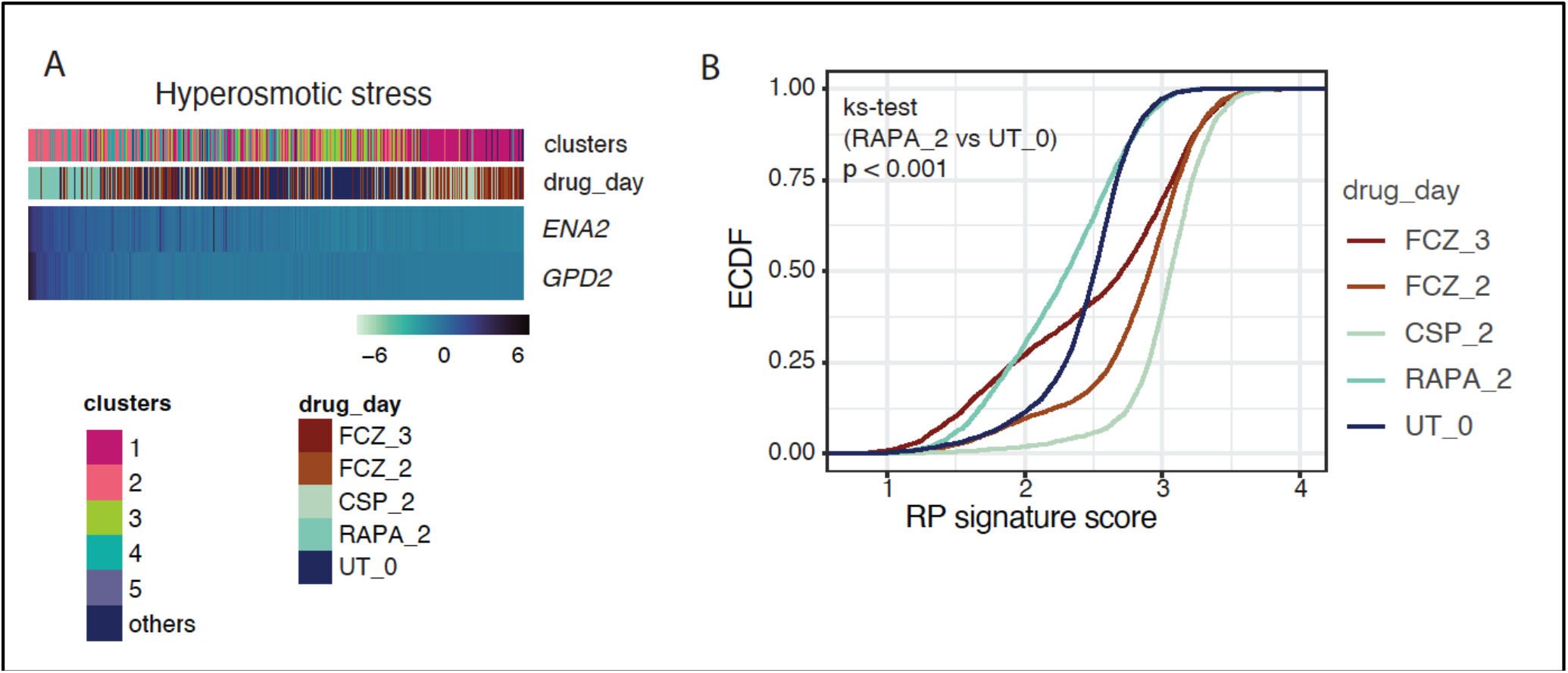
**(A)** The heatmap depicts genes (rows) in the hyperosmotic stress signature with significant variability and consistent expression across cells (columns). We used MAGIC to impute expression levels (z-score, color bar). Cells are linearly ordered by the overall magnitude of gene expression (mean gene rank). “clusters” (top) indicates the Leiden cluster of origin for each cell and “drug_day” indicates the condition for each cell. (**B**) Empirical cumulative distribution (ECDF) of signature scores for each cell under the different conditions (color) (ks-test, Kolmogorov-Smirnoff test).

## Notes

### Competing Interest Statement

The authors have declared no competing interest.

